# Forest canopy height change due to demographic and allometric shifts

**DOI:** 10.64898/2026.09.30.755816

**Authors:** Jeremy W. Lichstein, Michaela J. Woods, J. Aaron Hogan, Grant M. Domke

**Affiliations:** University of Florida, Department of Biology, Gainesville, Florida 32611, USA; Texas ACM University, Department of Ecology C Conservation Biology, College Station, Texas 77843, USA; Northern Research Station, United States Department of Agriculture, St. Paul, Minnesota, 55108, USA

**Keywords:** allometry, biomass, carbon storage, climate change, demography, drought, forest dynamics, Forest Inventory and Analysis program, forest monitoring, plant hydraulics

## Abstract

Forest canopy height is sensitive to shifts in demography (which affect tree size distributions and abundances of short-vs. tall-statured species) and allometry. Demographic and allometric effects on canopy height have not previously been studied across a large region. We analyzed more than two million individual tree measurements from about 60,000 remeasured inventory plots in non-plantation forests across 19 ecoregions of the conterminous United States to quantify changes in aboveground live tree biomass (AGB), canopy height, and its demographic and allometric components from ∼2003-2025. We tested the hydraulic limitation hypothesis, which predicts decreasing tree height with increasing hydraulic stress. AGB was strongly correlated with canopy height across ecoregions, and AGB change was strongly correlated with canopy height change over time. Canopy height change was positive in nearly all eastern ecoregions but varied in sign across western ecoregions. Demographic shifts led to increased canopy height in most eastern ecoregions but decreased canopy height in most western ecoregions. In contrast, allometric shifts, averaged across species, caused either positive or no significant change in ecoregion-mean canopy height. Our results were consistent with several predictions of the hydraulic limitation hypothesis: (*i*) intraspecific allometric shifts were more positive for understory than for canopy trees, (*ii*) across ecoregions, canopy height was positively correlated with precipitation and negatively correlated with vapor pressure deficit, and (*iii*) canopy height change over time was positively correlated with precipitation change. However, our results suggest modifications to the hydraulic limitation hypothesis, because allometric shifts tended to be positive, even in ecoregions with decreasing precipitation and increasing vapor pressure deficits. Such allometric shifts, which seem counter to the hydraulic limitation hypothesis, may be caused by CO_2_-driven increases in water-use efficiency. Whatever their cause, allometric shifts should be accounted for to avoid biased estimates of canopy height change and biomass change.

## Introduction

Forest canopy height affects diverse ecosystem and land-surface properties (Jucker et al., 2018; Migliavacca et al., 2021; Z. Zhang et al., 2022) and is inherently linked to forest biomass and carbon storage (Chave et al., 2019; Jucker et al., 2017; Lefsky et al., 2002). Canopy height can be directly estimated from remotely sensed data (Lang et al., 2023; Simard et al., 2011; Wagner et al., 2025), providing the potential for enhanced monitoring of biomass and other height-related forest properties (Chambers et al., 2007; Hurtt et al., 2004; Jucker et al., 2018; Travers-Smith et al., 2024). However, individual- and stand-level relationships between height, biomass, and other attributes may change over time due to shifts in species composition and allometry. Thus, understanding and accurately monitoring forest properties requires individual-tree-level observations of species identity and allometry that are absent from most remotely sensed data products, particularly those derived from widely available satellite data.

Forest demography—the tree-level growth, mortality, and recruitment rates that determine community dynamics—may affect canopy height through multiple pathways. Disturbance-recovery dynamics strongly affect tree size distributions and thus have a strong and direct effect on canopy height (Bormann & Likens, 1979; Peet, 1981). In addition, demography can also lead to shifts in species composition, which can affect canopy height due to species differences in allometry and maximum height. For example, forest maturation may involve shifts towards either taller- or shorter-statured tree species depending on environmental conditions and the traits of early- and late-successional species in a given region (Franklin & Hemstrom, 1981; Pare & Bergeron, 1995; Pastor et al., 1987; Van Cleve & Viereck, 1981). Furthermore, rising vapor pressure deficits (VPD) may select for shorter-statured species that are less susceptible to hydraulic failure (McDowell et al., 2020; McDowell & Allen, 2015).

In addition to demography, within-species shifts in height allometry—the non-linear relationship between individual height and stem diameter—may affect the height distribution of individual trees and, in turn, forest canopy height. Height allometries reflect the balance of the costs and benefits of increased height for a given diameter (Falster & Westoby, 2003). Costs may include hydraulic stress (Koch et al., 2004; McDowell & Allen, 2015; Ryan et al., 2006), allocational tradeoffs (Falster & Westoby, 2003), and increased exposure to wind damage (King, 1986; Lohmander & Helles, 1987; Thomas et al., 2015). Benefits may include increased light exposure relative to neighboring competitors (Falster & Westoby, 2003; Pacala et al., 1996) and wider dispersal of pollen and seeds (Nathan et al., 2001, 2002; Okubo & Levin, 1989). Increasing hydraulic stress should favor shorter allometries, whereas increases in stand biomass or density—e.g., due to forest maturation, fire suppression, or CO_2_ fertilization—should favor taller allometries (Forrester et al., 2017; Lines et al., 2012). In a free-air CO_2_ enrichment experiment, shifting height allometries accounted for 22% of the biomass increase in high CO_2_ plots (Kim et al., 2020). This CO_2_ effect may include direct effects related to increased water-use efficiency and reduced hydraulic stress (Keenan et al., 2013; van der Sleen et al., 2015), as well as indirect effects due to stand structure and light competition (Berntson & Wayne, 2000; Forrester et al., 2017; Lines et al., 2012).

Global trends toward rising VPD and intensifying disturbance regimes suggest that forest canopy heights should be decreasing in many regions (McDowell et al., 2020), but empirical tests of this hypothesis are lacking. Forests of the conterminous United States (U.S.) provide a valuable opportunity to study canopy height change because these forests span a wide range of climates and disturbance regimes and because the U.S. Forest Service has conducted systematic, standardized inventories of these forests over multiple decades, including observations of the stem diameters and heights of millions of individual trees (Bechtold & Patterson, 2005; Tinkham et al., 2018).

Here, we use data from the U.S. nationwide forest inventory (NFI) to quantify changes in canopy height and its demographic and allometric components across the conterminous U.S. from ∼2003-2025. Our study is based on measurements of individual trees in forest inventory plots, allowing us to quantify changes in demography (including shifts in diameter distributions and species composition) and within-species shifts in allometry. Our study is complementary to remote-sensing-based studies of canopy height, which have greater potential for spatiotemporal coverage but do not provide the tree-level species and diameter data needed to decompose height change into demographic and allometric components. We address the following questions: (*i*) How are aboveground biomass, canopy height, and their changes over time related across U.S. ecoregions? (*ii*) How have demographic and allometric shifts contributed to canopy height change? (*iii*) How is canopy height related to climate variables (temperature, precipitation, and VPD), and how is canopy height change (and its components) related to climate change? (*iv*) Do allometric shifts differ between canopy and understory trees?

We evaluate our results in the context of the hydraulic limitation (HL) hypothesis, which posits that tree heights are limited by hydraulic stress (Koch et al., 2004; McDowell & Allen, 2015; Moles et al., 2009; Ryan et al., 2006; Ryan & Yoder, 1997; Tao et al., 2016). We define the HL hypothesis broadly to encompass growth, mortality, and allometry, with the general prediction that tree heights should decrease as hydraulic stress increases (e.g., due to mortality of tall trees; McDowell & Allen, 2015). The HL hypothesis is supported by some studies of tree physiology (e.g., Koch et al., 2004; Ryan et al., 2006) and geographic patterns of canopy height (Moles et al., 2009; Tao et al., 2016). Here, we present novel tests of the HL hypothesis in terms of (*i*) changes over time in forest canopy height (and its demographic and allometric components) in relation to climate change, and (*ii*) how canopy and understory allometries shift over time within species. We assume that, on average, canopy trees experience greater hydraulic stress than understory trees (Myers et al., 1987; Niinemets, 2007). Thus, the HL hypothesis predicts that allometric shifts for understory trees will be more positive (or less negative) than for canopy trees.

## Methods

### Overview

We used data from ∼60,000 remeasured NFI plots (∼2 million measurements of individual trees) in non-plantation forests across 19 ecoregions of the conterminous U.S. from ∼2003-2025 to (*i*) quantify relationships between stand-level aboveground wood biomass (AGB), forest canopy height, and their changes over time; and (*ii*) partition changes in canopy height into demographic and allometric components. Our partitioning approach involved fitting static and dynamic height-diameter allometries for each tree species in each ecoregion and combining these models with raw forest inventory data. We evaluated correlations between height variables (observed change and modeled demographic and allometric components) and variables describing baseline climate conditions and climate change. To gain insights into potential drivers of allometric shifts, we compared allometric shifts between canopy and understory trees. Abbreviations and symbols used frequently in the text are defined in Table 1. Sample sizes are given in Table S1. All analyses were performed in R (R Core Team, 2025).

**Table 1.** Terms used frequently in the text. See Methods for details.

| Symbol | Definition |
| --- | --- |
| AGB | Aboveground wood biomass of live trees in an inventory plot. Ecoregion values are means across plots. |
| Canopy tree | Live tree in an inventory plot with an open-grown, dominant, or codominant FIA crown class. This designation is based on relative, rather than absolute, height and is equivalent to “sun exposed.” |
| DBH | Stem diameter of an individual tree reported by FIA. For simplicity, we use “DBH” (diameter at breast height) to refer to diameters of all trees, including woodland species whose diameters are measured at a point lower than at breast height. |
| FIA | Forest Inventory and Analysis program and database |
| $h$ | Height of an individual tree reported by FIA, including any effects of damage or dieback. |
| $H_{\max}$ | Maximum height of live trees in an inventory plot. Ecoregion values are means across plots. |
| $H_{\text{mean}}$ | Mean height of live canopy trees in an inventory plot. Ecoregion values are means across plots. |
| NFI | U.S. nationwide forest inventory conducted by the FIA program |
| No-management dataset | Non-plantation inventory plots with no reported management (e.g., harvest) between the initial and final plot measurements in our analysis dataset. These plots may have been managed prior to our study. Results we report are for the no-management dataset unless stated otherwise. |
| Non-plantation forests | Plots classified by FIA as natural stands (no clear evidence of artificial regeneration), which does not imply an absence of management; e.g., non-plantation forests may be harvested. |
| $t = 0$ | Initial measurement date of an inventory plot. We use “$t = 0$” to refer collectively to the initial measurements of all plots; in reality, each plot has a different measurement date. |
| $t = T$ | Final measurement date of an inventory plot. We use “$t = T$” to refer collectively to the final measurements of all plots; in reality, each plot has a different measurement date. |
| VPD | Vapor pressure deficit of the atmosphere. |
| $\Delta H_{\text{allom}}$ | Allometry component of height change, which represents height change due to intraspecific shifts in allometry, holding the DBH distribution and species composition constant. Ecoregion values are means across plots. |
| $\Delta H_{\text{dem}}$ | Demography component of height change, which represents height change due to shifts in both the DBH distribution and species composition, holding intraspecific allometries constant. Ecoregion values are means across plots. |

### Forest inventory data

The NFI is one of several inventories conducted by the Forest Inventory and Analysis (FIA) program. For simplicity, we hereafter use “FIA” to refer to the NFI design and data. We downloaded the FIA data (Burrill et al., 2025) on 7 April 2026 from https://research.fs.usda.gov/products/dataandtools/fia-datamart. We restricted our analysis to plots that were measured and remeasured using FIA’s national standardized design (summarized below), which was implemented starting in the early 2000s in most U.S. states (Bechtold & Patterson, 2005) and includes diameter and height measurements of individual trees (Burrill et al., 2025). For our study, we selected a subset of the national standardized data to create a consistent dataset across the conterminous U.S., with each plot location represented by two plot measurements (initial and final), separated by 9-13 years, that met our filtering criteria (described below). The 9-13-year range maximized the number of remeasured plots available for our analysis. Since ∼2000, western U.S. FIA plots have been remeasured every ∼10 years, and eastern U.S. FIA plots have been remeasured every ∼5 years. Therefore, in the eastern U.S., we typically skipped an intermediate plot measurement to achieve the target 9-13-year interval. In cases where multiple 9-13-year intervals were available for a given plot location (which was common in the eastern U.S. but rare in the western U.S.), we only used the most recent interval, so that each plot in our analysis was represented by two measurements. Rather than using all available data, our approach prioritized consistency to facilitate straightforward comparisons across different U.S. regions. Across all plots in our study, the mean initial and final dates were ∼2009 and ∼2020, respectively, with a mean difference of 10.7 years. The temporal coverage of the dataset was ∼22 years: the 2.5^th^ percentile of initial measurement dates was ∼2003, and the 97.5^th^ percentile of final measurement dates was ∼2025.

Each FIA plot includes four subplots, distributed across 0.4 ha. Trees with a diameter ≥ 12.7 cm are tallied in circular subplots of radius 7.32 m; in some western U.S. plots, this radius is expanded to 17.95 m for large diameter trees (e.g., diameter ≥ 61 or 76 cm, depending on local protocols). Each subplot includes a microplot (2.07 m radius) for tallying trees with trees with diameter ≥ 2.54 cm. Stem diameter is measured for all live tally trees. For most tree species, FIA measures stem diameter at 1.37 m height (i.e., diameter at breast height, DBH). For woodland tree species, which are often multi-stemmed, FIA measures the diameter of each stem at the root collar or ground line and reports the square root of the sum of squared diameters. For conciseness, we hereafter refer to the diameter of all trees as “DBH.” Other key variables reported for tally trees include species, status (live or dead), mortality agent for dead trees (fire, insect, harvest, unknown, etc.), total and actual height (for undamaged trees, these two height measures are equal; see details below), crown class (overtopped, intermediate, codominant, dominant, or open grown), aboveground wood biomass (AGB), and the number of trees per-unit ground area that the tally tree represents (the inverse of its DBH-dependent plot sampling area). For each tally tree, FIA reports AGB estimated from species- and region-specific biomass allometries that incorporate field observations of DBH, height, and rotten or missing wood (Westfall et al., 2024).

Each FIA plot measurement includes one or more mapped “conditions” defined by land ownership, stand origin (natural stands vs. those with clear evidence of artificial regeneration), disturbance history, forest type, soil type, and stand structure. We restricted our analysis to field-sampled, non-plantation, forest land conditions. We defined non-plantation conditions as “natural stands” according to FIA’s stand origin code. FIA defines “forest land” as having ≥ 10% tree canopy cover at the time of sampling or in the past (based on the presence of snags or stumps) in an area at least 0.4 ha in size and 36.6 m wide. We restricted our analysis to plot measurements in which at least 95% of the plot area remained after excluding non-sampled, plantation (artificial regeneration), and non-forest conditions.

We analyzed two FIA datasets: (*i*) all non-plantation forests, and (*ii*) non-plantation forests with no reported evidence of management activities between the initial and final measurements of each plot (“no-management dataset”). Despite being classified by FIA as “natural stands,” management activities may still occur in non-plantation plots, including clearcut or selective harvest. We considered a plot to be managed if, between its initial and final measurements, FIA reported any harvested trees in the FIA Tree table or any treatments (other than “natural regeneration”) in the FIA Condition table. Of the plots where no management was reported, many were likely managed in prior decades (Canham et al., 2013). Thus, our no-management dataset does not rule out indirect effects from prior management. Throughout this paper, sample sizes and results are for the no-management dataset unless stated otherwise. Sample sizes by ecoregion for both datasets are provided in Table S1.

We assigned each plot to one of 19 ecoregions (Fig. S1) by aggregating the ecological subsection codes reported by FIA to the division level (McNab et al., 2007). Each ecoregion represents an area of similar climate and broad vegetation type. We used the division level because a recent analysis of FIA data found that the finer province level often had insufficient sample size to quantify shifts in forest dynamics (Hogan et al., 2024). We mapped our results to ecoregions using shapefile boundaries accessed from https://data.fs.usda.gov/geodata/edw/datasets.php on 22 September 2024.

We excluded palm trees from our analysis because they lack secondary diameter growth (Goodman et al., 2013) and thus do not have meaningful height-diameter relationships. To avoid artifacts caused by excluding palms (e.g., underestimating canopy heights of plots dominated by palms), we only included plots with palms if there were at least four live canopy trees at both the initial and final plot measurements, and if palms comprised no more than 25% of the canopy trees at either measurement. This ensured that for plots with palms, at least three non-palm trees remained at both measurements after excluding palms. For non-palm plots, we imposed no such constraints; e.g., we included non-palm plots in our analysis even if no live canopy trees were present, as long as the plots met our other filtering criteria. All palms that were removed belonged to a single species (*Sabal palmetto*) in the southeast U.S. (ecoregions 230 and 410; Fig. S1). Palm-filtering removed 0.5% of canopy trees from ecoregion 230 and 6.4% of canopy trees from ecoregion 410.

### Defining canopy and understory trees

To identify canopy trees for quantifying forest canopy height and to distinguish canopy (sun-exposed) from understory (shaded) trees for tests of the hydraulic limitation hypothesis, we assigned FIA tally trees to canopy and understory strata based on the FIA crown class reported at a given measurement time (e.g., a given tree could be an understory tree at the initial plot measurement and a canopy tree at the final measurement). We defined “canopy trees” as having an open-grown, dominant, or codominant crown-class, and “understory trees” as having an overtopped crown-class. Trees with an intermediate crown-class were included in analyses based on all trees (e.g., plot-level AGB and total tree density) but were excluded from analyses focused on canopy or understory trees. Note that FIA crown classes are based on relative, not absolute, height. Thus, “canopy trees” in our analysis are sun-exposed from above. In low-density stands with no shaded trees, all individuals would qualify as “canopy trees,” regardless of their height.

### Observed changes in biomass and tree density

To quantify observed ecoregion-level patterns in AGB and its change, we first calculated AGB (Mg ha^−1^) for each plot at time *t* (initial and final) as *AGB_t_* = ∑*_i_ b_i_ n_i_*, where the sum is over all live trees at time *t*, *b_i_* is AGB (Mg) for tree *i* at time *t*, and *n_i_* is the number of trees per hectare that the tree represents at time *t* (TPA_UNADJ in the FIA Tree table, converted to ha^−1^ units). Plots with no live trees at time *t* have, by definition, *AGB_t_* = 0. We calculated AGB change (Mg ha^−1^ yr^−1^) for each plot as Δ*AGB* = (*AGB_T_* − *AGB*_0_)/*T*, where *T* is the number of years between the initial (*t* = 0) and final (*t* = *T*) measurements. To quantify broad-scale patterns in AGB and its change, we averaged *AGB*_0_ and Δ*AGB* across plots in each ecoregion. We calculated relative AGB change (% yr^−1^) for each ecoregion as 100 · Δ*AGB* / *AGB*_0_ (here, the symbols Δ*AGB* and *AGB*_0_ refer to ecoregion means). We did not quantify relative change at the plot level because some plots had *AGB*_0_ = 0.

As in the analysis described above for AGB, we quantified observed ecoregion-level patterns in live tree density and its change. We first calculated live tree density (trees ha^−1^) for each plot at time *t* as *N_t_* = ∑*_i_ n_i_*, where *n_i_* is defined above. We then calculated Δ*N* for each plot, and ecoregion-level means of *N*_0_ (trees ha^−1^), Δ*N* (trees ha^−1^ yr^−1^), and relative Δ*N* (% yr^−1^), as explained for AGB above. We implemented two versions of the tree density analysis: for all live trees and for live canopy trees.

### Observed changes in canopy height

We quantified observed ecoregion-level patterns in canopy height and its change based on two different plot-level definitions of canopy height: (1) the mean height of live canopy trees in a plot (*H*_mean_); and (2) the height of the tallest live tree in a plot (*H*_max_). Plots with no live trees at a given time, by definition, have *H*_mean_ = *H*_max_ = 0. We estimated ecoregion-level patterns for *H*_mean_ and *H*_max_ in a similar manner as for AGB above: we weighted each canopy tree by its trees-per-hectare value when calculating *H*_mean_; we calculated plot-level *H*_mean_, *H*_max_, and their changes (Δ*H*; m yr^−1^) from *t* = 0 to *t* = *T*; we averaged the *t* = 0 and change values across plots within each ecoregion for both definitions of canopy height (*H*_mean_ and *H*_max_); and we calculated ecoregion-mean percent change (% yr^−1^) in *H*_mean_ and *H*_max_ relative to their ecoregion means at *t* = 0. We estimated the width of 95% confidence intervals (CIs) for observed ecoregion mean change as ± 1.96 × s.e., where s.e. is the standard error of the mean.

In the height analysis described above, we defined individual tree height as the “actual height” reported by FIA (ACTUALHT in the FIA Tree table), which includes the effects of any damage or dieback (Burrill et al., 2025). FIA defines actual height as the length (to the nearest 0.3 m) from the ground along the bole to the tip of the apical meristem for an undamaged crown, to the highest living portion of a recently damaged crown, and to the tip of a qualifying leader for a damaged and regrowing crown (see “actual length” in USDA, 2022). For each tree, FIA reports a code indicating if height was field measured, visually estimated, or modeled (HTCD in the FIA Tree table). To quantify observed patterns in canopy height and its change, we did not filter on this code, because we aimed to minimize missing data that could bias our results. For example, if the heights of the tallest trees were visually estimated or modeled, excluding these trees would cause negative bias in *H*_mean_ and *H*_max_. Similarly, if all tree heights in a plot were visually estimated or modeled, excluding all these heights would yield erroneous zero values for *H*_mean_ and *H*_max_. We expect errors associated with visually estimated or modeled heights to have little effect on our ecoregion-level estimates of observed canopy height and its change, because 90% of canopy tree heights in the FIA dataset we analyzed were field measured.

### Allometry models

Our method of partitioning height change into demographic and allometric components (described in a later section) requires both static and dynamic height allometries for each species in each ecoregion. Static models describe how individual height (*h*) depends on DBH at the time of the initial plot measurements, and dynamic models describe how allometries change between the initial (*t* = 0) and final (*t* = *T*) plot measurements. We defined *h* as the “actual height” reported by FIA (see details above). We restricted the allometry fitting dataset to live trees with field-measured *h* (according to HTCD in the FIA Tree table) in the no-management dataset (Table 1).

Within each ecoregion, we fit separate models for each species × ecoregion × canopy position (canopy or understory) with at least 30 observations in both the *t* = 0 and *t* = *T* datasets. This yielded 642 canopy allometries and 418 understory allometries. We used understory allometries only to compare allometric shifts between canopy and understory trees (see details in a later section). Unless stated otherwise, all analyses in our paper are based on canopy allometries.

We fit allometries using nonlinear least squares with the *nlsLM* function in the *minpack.lm* R package (Elzhov et al., 2023). Due to the impracticality of fitting > 1000 nonlinear mixed models (canopy and understory cases combined), we did not include individual-level random effects in the dynamic models (which have repeat observations for some trees; see below) or plot-level random effects in either the static or dynamic models. Our allometric parameter estimates should nevertheless be approximately unbiased, but their standard errors may be biased (Liang & Zeger, 1986). Our main conclusions are based only on the parameter estimates.

For the static models, we quantified the relationship between *h* (m) and DBH (cm) using an asymptotic function (Pacala et al., 1994; Thomas, 1996):

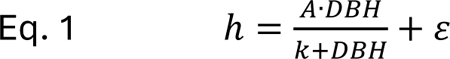

where *A* (m) is the asymptote, *k* (cm) is the DBH at which the expected height is *A*/2, and *ε* is a normally distributed error. We fit static allometries (Eq. 1) using only the *t* = 0 data from each plot.

For the dynamic models, we assumed that the asymptote changed linearly over time:

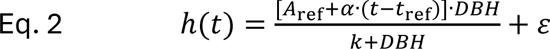

where *A*_ref_ (m) is the asymptote in the reference year, *α* is the rate of change in the asymptote (m yr^−1^), *t*_ref_ is the reference year, the term in square brackets is the asymptote in year *t*, and other terms are as in Eq. 1. If *α* > 0, the *h* vs. DBH curve shifts up over time (i.e., *h* increases with *t* for a given DBH); if *α* < 0, the curve shifts down over time; and if *α* = 0, Eq. 2 is identical to Eq. 1. The choice of *t*_ref_ affects *A*_ref_ but does not affect *α* or any conclusions drawn from Eq. 2. We arbitrarily assigned *t*_ref_ = 2000. We fit dynamic allometries (Eq. 2) using both the *t* = 0 and *t* = *T* data from each plot. An individual tree could be included zero, one, or two times in the fitting dataset for a given dynamic model (canopy or understory) depending on its canopy position and whether it was alive with a field-measured height at *t* = 0 and *t* = *T*.

We evaluated the functional forms (Eqs. 1-2) by visually inspecting (*i*) fitted curves of *h* vs. DBH overlaid on scatterplots of the raw data (left columns of Figs. S2-S3) and (*ii*) plots of observed vs. predicted *h* for data aggregated into bins (right columns of Figs. S2-S3). The goal of binning was to minimize the effects of measurement error and individual variation when evaluating the functional forms. We created bins as follows: For each model, we assigned the *n* observations to DBH bins of equal sample size (or nearly equal, if *n* was not evenly divisible by the number of bins). If *n* ≤ 300, we created 10 bins. If *n* > 300, we assigned at least 30 observations per bin, and we created up to 30 bins. For example, if *n* = 600, we created 20 bins of 30 observations each; or if *n* = 1500, we created 30 bins of 50 observations each. The individual- and bin-level visual evaluations indicated good fits in nearly all cases. We also explored a modification of Eqs. 1-2 in which DBH was raised to a fitted exponent, but this additional parameter rarely provided a significant improvement in model fit and was not included in our results.

We further evaluated the functional forms by calculating the proportion of variance explained, 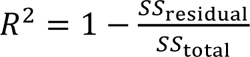, at individual and bin levels for each model (Fig. S4). At the individual level (*R*^2^_ind_), we calculated *SS*_residual_ as the sum of squared model residuals, and *SS*_total_ as the sum of squared mean-centered *h* values. Mean *R*^2^_ind_ for both static and dynamic models were > 0.5 (Fig. S4a-b). At the bin level (*R*^2^_ind_), we calculated observed and predicted mean *h* values within each bin (*h*_bin_). We then calculated *SS*_residual_ as the sum of squared differences between observed and predicted *h*_bin_, and *SS*_total_ as the sum of squared differences between each observed *h*_bin_ and the mean observed *h*_bin_. Because much of the individual noise is removed at the bin level (especially for large *n*), and because *R*^2^_ind_ penalizes for departures from the 1:1 line, we considered *R*^2^_ind_ to be a useful metric for evaluating our functional forms. Most *R*^2^_ind_ were > 0.9, with most of the lower *R*^2^_ind_ values occurring for models with small *n* (Fig. S4c-d). The small-*n* cases have minimal impact on our main results and conclusions, because the effect of each allometric model on our overall results scales with its *n*.

### Partitioning canopy height change into demography and allometry components

We partitioned canopy height change into two components: allometry and demography, where “demography” includes shifts in the DBH distribution (e.g., due to disturbance or forest maturation), as well as shifts in species composition (e.g., shifts from short- to tall-allometry species, or vice versa). We implemented this analysis for both definitions of canopy height (*H*_mean_ and *H*_max_). The demography component represents height change resulting from shifts in both the DBH distribution and species composition, holding intraspecific allometries constant. Conversely, the allometry component represents height change due to intraspecific shifts in allometry, holding the DBH distribution and species composition constant. We also explored a more complex method that partitioned the demography component into separate DBH and species components, which yielded a poorer fit to observations than the simpler method with a single combined demography component. Results in this paper are based on a single combined demography component.

For each plot, we estimated the demography component of height change, Δ*H*_dem_ (m yr^−1^), by using static allometries (Eq. 1) to predict *h* for each live tree at *t* = 0 and *t* = *T* based on its observed DBH and species identity at *t* = 0 and *t* = *T*. Because we fit static allometries using only the *t* = 0 data, these predictions control for any intraspecific shifts in allometry between *t* = 0 and *t* = *T*. For individuals of species that did not meet our sample-size criteria for model-fitting in a given ecoregion (2.6% of individuals), we used the static allometry for the ecoregion’s species with the largest sample size. Then, using the procedures previously described for observed height change, we used the predicted *h* values to calculate plot-level *H*_mean_, *H*_max_, and their changes (Δ*H*_dem_) from *t* = 0 to *t* = *T*. We then averaged over the plot-level values to calculate ecoregion means of Δ*H*_dem_for *H*_mean_ and *H*_max_.

For each plot, we estimated the allometry component of height change, Δ*H*_allom_(m yr^−1^), by using the dynamic allometries (Eq. 2) to predict *h* at *t* = 0 and *t* = *T* for each live tree present at *t* = 0, holding DBH and species identity constant at the observed *t* = 0 values. Thus, we predicted how *h* values would have changed in each plot from *t* = 0 to *t* = *T* if there were no changes in DBH or species composition and no mortality or recruitment. This isolates the effects of intraspecific shifts in allometry (*α* in Eq. 2). For individuals of species that did not meet our sample-size criteria for model-fitting in a given ecoregion (2.6% of individuals), we used the dynamic allometry for the ecoregion’s species with the largest sample size. We then used the predicted *h* values to calculate plot-level Δ*H*_allom_ and ecoregion means of Δ*H*_allom_for *H*_mean_ and *H*_max_, as described for Δ*H*_dem_above.

For each plot, we predicted total height change, Δ*H*′, as the sum of the demography and allometry components: Δ*H*′ = Δ*H*_dem_ + Δ*H*_allom_. We then averaged the plot-level predictions to calculate the ecoregion mean of Δ*H*′ for both *H*_mean_ and *H*_max_. Multiple factors may contribute to mismatches between predicted (Δ*H*′) and observed (Δ*H*) ecoregion-mean height-change, including spatial variation in allometry within ecoregions and interactions (covariance) between the components (Δ*H*_dem_ and Δ*H*_allom_). For example, if species that gain in abundance from *t* = 0 to *t* = *T* tend to have more positive (or less negative) intraspecific allometry shifts, then the interaction term will be positive, and ignoring it will lead to underpredicting observed change. In contrast, if species that gain in abundance tend to have less positive (or more negative) intraspecific allometry shifts, then the interaction term will be negative, and ignoring it will lead to overpredicting observed change. For simplicity, we ignored the interaction terms. The close match between Δ*H*′ and Δ*H* (see Results) suggests that interactions or other sources of error are, on average, small in our analysis.

To quantify uncertainty in the six ecoregion means (*H*_mean_ and *H*_max_ for each of Δ*H*_dem_, Δ*H*_allom_, and Δ*H*′), we considered uncertainty due to sampling variance (plot-to-plot variation) and uncertainty in allometry parameter values. For each of the six means, we estimated its sampling variance (*σ^2^_samp_*) as the square of the standard error of the mean across plots, evaluated at the best-fit allometry parameter values. For each of the six means, we estimated the variance due to parameter uncertainty (*σ^2^_par_*) as the variance in the mean across 1000 parameter samples. We generated these samples from the estimated sampling distributions of the allometry parameters (Eqs. 1-2), which we assumed to be multivariate normal (MVN), with the MVN mean given by the best-fit parameter values and covariance matrix extracted using the *vcov* function in R (Bolker, 2008). We rejected the very rare parameter samples with *k* < 0.1 (i.e., if the half-asymptotic DBH was less than 0.1 cm) because such values are unrealistic and because *k* < 0 can cause numerical artifacts in Eqs. 1-2. We assumed the two variance components were independent, and we combined them as follows: Let *θ* be one of the six ecoregion means (*H*_mean_ and *H*_max_ for each of Δ*H*_dem_, Δ*H*_allom_, and Δ*H*′). We estimated the variance of *θ* as 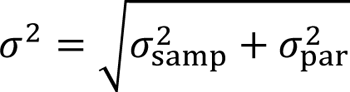. We report *θ* evaluated at the best-fit parameter values, *θ*^, and its 95% CI as *θ*^ ± 1.96 × *σ*.

### Climate data and correlations

We quantified correlations between canopy height and baseline climate, and between canopy height change (and its components) and climate change. In these analyses, all height and climate variables were represented as ecoregion-level means, because the allometry models used to estimate the height-change components were fit at the ecoregion level. To minimize the management effects, we used the no-management dataset for these analyses. For each ecoregion, we estimated 12 climate variables: baseline means (1990-2002) and trends (2003-2025) for temperature, precipitation, and VPD for both annual (12 months) and summer (June, July, August) time windows. For example, for each ecoregion, baseline annual temperature is the mean annual temperature from 1990-2002, baseline summer temperature is the mean summer temperature from 1990-2002, and the annual and summer temperature trends are estimated from 2003-2025. The trend period spans the 2.5^th^ and 97.5^th^ percentiles of the measurement dates in the FIA data we analyzed.

We estimated the above climate variables using the PRISM 4-km gridded monthly weather data (PRISM Group, 2026). We extracted the PRISM data for each FIA plot location in our analysis based on the approximate spatial coordinates reported by FIA, which are within 1.6 km of the true locations for most plots (Burrill et al., 2025). Thus, the ecoregion-level mean climate variables we calculated are representative, on average, of the plots included in our analysis. Given the ecoregion scale of our climate analysis, we would not expect the resolution of the FIA plot coordinates or the PRISM data to affect our results. For each plot, we calculated the baseline mean (1990-2002) and linear trend (2003-2025), and then we averaged the baseline means and trends across plot locations within each ecoregion.

We calculated Pearson’s correlation (*r*) at the ecoregion level (*n* = 19) for the following cases: For canopy height vs. baseline climate, we considered both definitions of canopy height (*H*_mean_ and *H*_max_) combined with the six baseline climate variables(annual and summer for each of temperature, precipitation, and VPD), resulting in 12 correlation tests. For *H*_mean_ and *H*_max_, we used their ecoregion-mean initial values (*t* = 0). For canopy height change vs. climate change, we considered six height-change variables (observed Δ*H* and the modeled components, Δ*H*_dem_and Δ*H*_allom_, for each of *H*_mean_ and *H*_max_) combined with the six climate trends variables (annual and summer for each of temperature, precipitation, and VPD), resulting in 36 correlation tests.

### Intraspecific allometric shifts

We used the dynamic allometry models (Eq. 2) to quantify intraspecific allometric shifts over time at a given reference DBH. This approach provided a measure of allometric shift that was easier to interpret than the rate of asymptotic change (*α* in Eq. 2), because in many models, *h* did not approach its asymptote within the observed DBH range (Figs. S2-S3). We defined the allometric shift, *δ* (m yr^−1^), as the rate of change in *h* for a tree of DBH *x*:

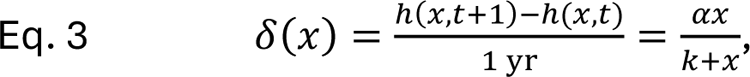

where the ℎ(·) terms are Eq. 2 evaluated with DBH = *x* at times *t* + 1 and *t*, and *α* and *k* are as defined in Eq. 1. The sign of *δ*(*x*) equals the sign of *α* (assuming *k* > 0). Thus, the reference DBH (*x*) affects the magnitude but not the sign of *δ*. For understory trees, we report allometric shifts using *x* = 10 cm. For canopy trees, we report allometric shifts using both *x* = 10 cm (for direct comparison with understory trees) and *x* = 30 cm (a typical DBH for mid-successional canopy trees in many U.S. forests). To quantify uncertainty in *δ*, we generated 1000 samples from the distribution of *α* and *k* (assumed to be MVN), and we used these parameter samples to generate a sampling distribution for *δ* (Bolker, 2008). We used these sampling distributions to estimate the variances described below. We rejected parameter samples with *k* < 0.1, as explained above.

We compared canopy and understory allometric shifts at *x* = 10 cm using paired *t*-tests of the difference in *δ* between canopy and understory trees of the same species × ecoregion. We performed separate *t*-tests for canopy-understory pairs in the eastern and western U.S. In the *t*-tests, we accounted for uncertainty in *δ* by weighting each difference by its inverse variance (the sum of the canopy and understory variances of *δ*, which we estimated from their sampling distributions). We tested the weighted mean difference against a null hypothesis of zero using a one-sample *t*-test.

## Results

### Patterns of biomass and canopy height change

AGB, canopy height (both *H*_mean_ and *H*_max_; see definitions in Table 1), and their changes over time were strongly positive correlated across U.S. ecoregions (Fig. 1 and Figs. S5-S7). Correlations between AGB (and its change) and *H*_max_ (and its change) were especially strong (Figs. S6-S7). In contrast, AGB and its change had weak relationships with canopy tree density, total tree density, and their changes (Figs. S6-S9). AGB and canopy height were relatively high in the East and along the Pacific coast, variable in the Interior West, and low in the Southwest (left columns of Fig. 1 and Fig. S5). Changes in AGB and canopy height tended to be positive in the East, varied in sign across the West, and were more positive (or less negative) in the no-management dataset (Fig. 1) compared to all non-plantation FIA plots (including those that were managed during our study; Fig. S5). Decreases in AGB and canopy height in the West were more apparent for relative change (% yr^−1^; right columns of Fig. 1 and Fig. S5) than for absolute change (m yr^−1^; middle columns of Fig. 1 and Fig. S5).

**Fig. 1.**
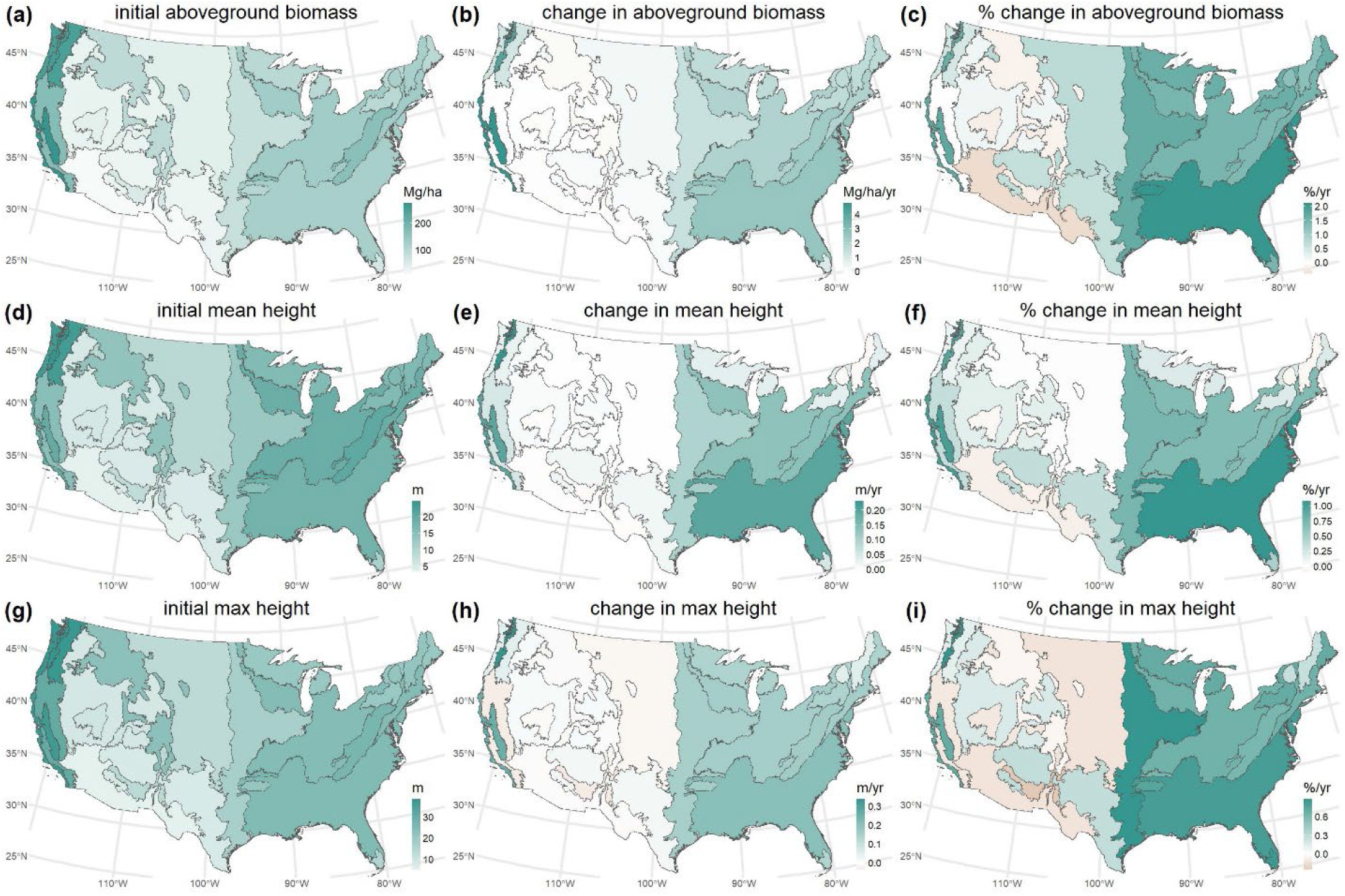
Changes in aboveground biomass **(a-c)**, mean canopy height **(d-f)**, and maximum tree height **(g-i)** for the no-management dataset (non-plantation inventory plots with no reported management activities during the study period). Colors indicate the mean value across inventory plots in each ecoregion (Fig. S1). **Left column:** initial measurement of each plot (mean initial date ∼2009). **Middle column:** change from the initial to final measurement of each plot (mean final date ∼2020). **Right column:** relative change = 100 × (middle column)/(left column).

### Demographic and allometric components of canopy height change

Predicted changes in ecoregion mean canopy height (the sum of the demography and allometry components, Δ*H*_dem_and Δ*H*_allom_) were strongly correlated along the 1:1 line with observed changes in *H*_mean_ and *H*_max_ (Fig. 2). Observed height change was more strongly correlated with Δ*H*_dem_ than with Δ*H*_allom_, but was typically underpredicted by Δ*H*_dem_alone because Δ*H*_allom_was typically positive (Fig. 2). Geographic patterns in Δ*H*_dem_differed somewhat between *H*_mean_ and *H*_max_; nevertheless, Δ*H*_dem_tended to be positive in the East and variable in the West for both *H*_mean_ and *H*_max_ (Fig. 3 and Figs. S10-S11). Geographic patterns in Δ*H*_allom_ were similar for *H*_mean_ and *H*_max_, tending to be positive in the East and positive or near-zero in the West (Fig. 3 and Figs. S10-S11). Δ*H*_allom_was not significantly negative for any ecoregion (Fig. 2b and 2e). On average, the magnitude of Δ*H*_dem_was roughly twice that of Δ*H*_allom_(Fig. S11), but this discrepancy was partly due to two western ecoregions (240 and 260) that are relatively small in area (Fig. S1). Thus, on an area basis, the difference in magnitude between Δ*H*_dem_ and Δ*H*_allom_ was more modest.

**Fig. 2.**
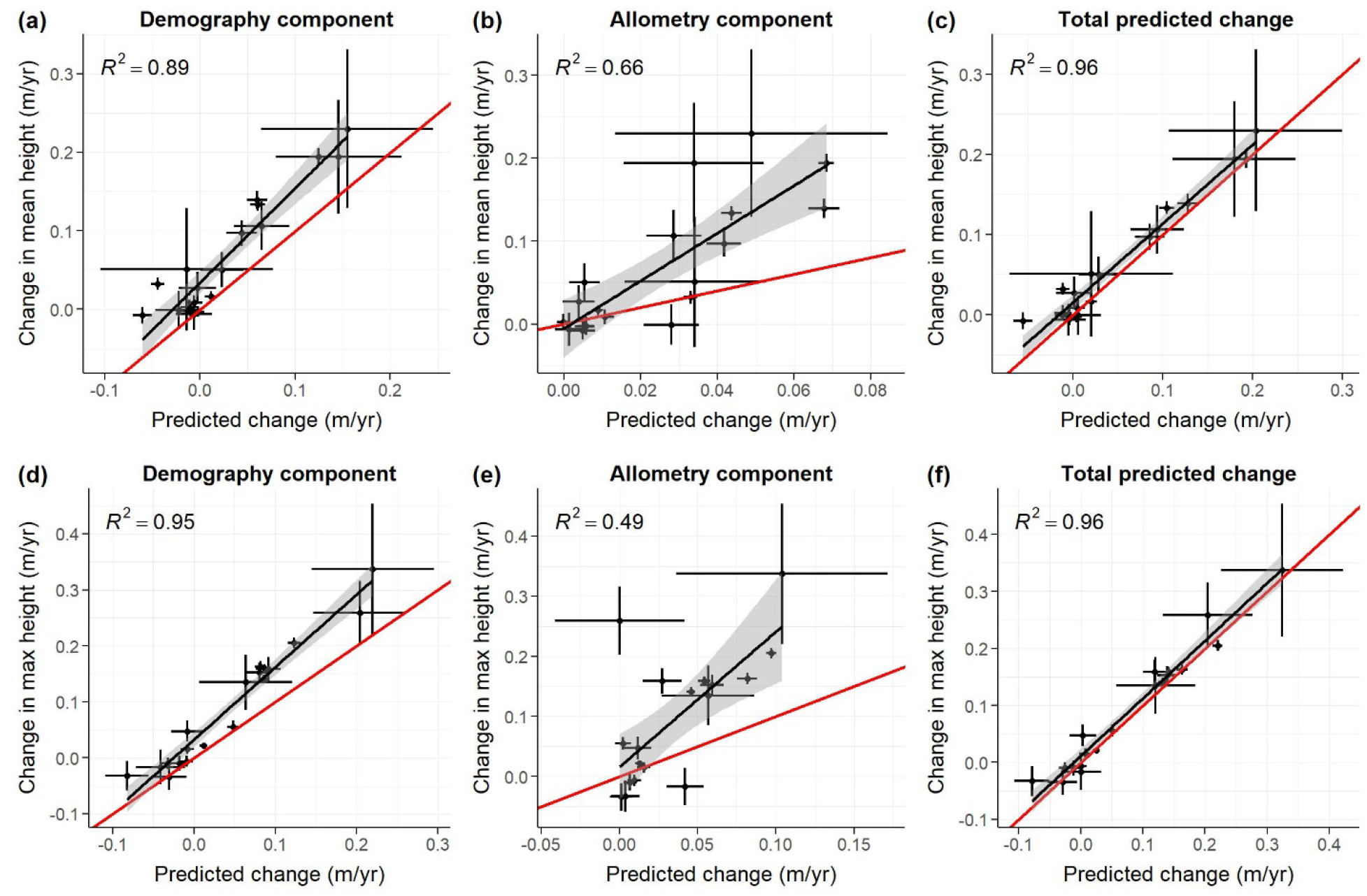
Observed vs. predicted change in mean canopy height **(a-c)** and maximum tree height **(d-f)** across U.S. ecoregions (*n* = 19) for the no-management dataset (non-plantation inventory plots with no reported management activities during the study period). Observed changes (y-axis) are ecoregion means (mean canopy height: Fig. 1e; max height: Fig. 1h) with 95% confidence intervals (CIs). Predicted changes (x-axis) are ecoregion means (Fig. S10) for demography and allometry components and their sum (total predicted change) with 95% CIs. Red lines are the 1:1 line. Black lines (with shaded 95% CIs) and *R*^2^ values are for regressions of observed vs. predicted change (all regressions have *p* < 0.001).

**Fig. 3.**
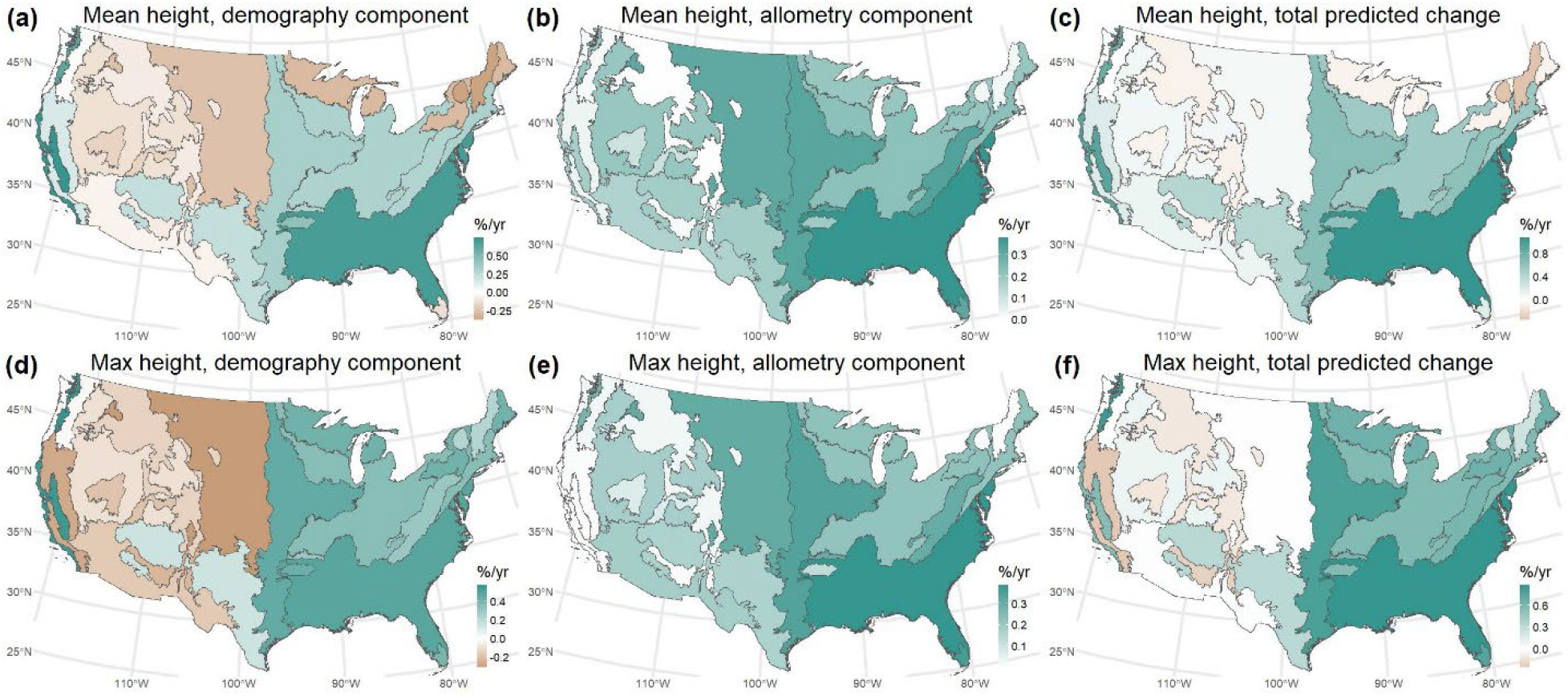
Predicted relative change (% yr^−1^) in mean canopy height **(a-c)** and maximum tree height **(d-f)** for the no-management dataset (non-plantation inventory plots with no reported management activities during the study period). Colors indicate the mean value across inventory plots in each ecoregion (Fig. S1). **Left column:** the demography component represents height change due to shifts in both tree diameter distributions and species composition, holding intraspecific allometries constant. **Middle column:** the allometry component represents height change due to intraspecific shifts in allometry, holding tree diameter distributions and species composition constant. **Right column:** total predicted change is the sum of the demography and allometry components. Predicted percent change is relative to initial observed ecoregion-mean canopy height (Fig. 1d) or maximum tree height (Fig. 1g).

### Relationships with climate variables

Correlations across ecoregions between canopy height and baseline climate (Fig. S12) and between canopy height change and climate change (Fig. S13) were either non-significant or were consistent with the hydraulic limitation hypothesis. Observed *H*_mean_ and *H*_max_ were positively correlated with annual precipitation and negatively correlated with annual and summer VPD (Table 2). Observed canopy height change and Δ*H*_allom_ were positively correlated with annual precipitation change for all four cases considered (*H*_mean_ and *H*_max_ expressed in m yr^−1^ and % yr^−1^ units; Table 2). In contrast, Δ*H*_dem_was correlated with annual precipitation change in only one of the four cases (*H*_max_ with % yr^−1^ units; Table 2). Annual precipitation change was the only climate change variable that was significantly correlated with any height change variable.

**Table 2.** Correlations across ecoregions (*n* = 19) between canopy height and baseline climate, and between canopy height change and climate change. Mean canopy height (*H*_mean_) and maximum tree height (*H*_max_) are ecoregion means for the initial measurement of each plot. Baseline climate variables (annual and summer for temperature, precipitation, and VPD) are ecoregion means over 1990-2002. Height change variables are ecoregion mean changes from the initial to final measurement of each plot and are analyzed with both absolute units (m yr^−1^) and relative to the initial ecoregion mean (% yr^−1^). Climate change variables (annual and summer for temperature, precipitation, and VPD) are mean ecoregion trends from 2003-2025. Pearson’s correlation (*r*) is reported only for significant cases: * *p* < 0.05, ** *p* < 0.01, *** *p* < 0.001.

| Height variable | Component | Climate variable | $r$ |
| --- | --- | --- | --- |
| Mean canopy height, $H_{\text{mean}}$ (m) | Initial observed | Baseline annual precipitation | 0.85*** |
|  |  | Baseline annual VPD | -0.74*** |
|  |  | Baseline summer VPD | -0.85*** |
| Maximum tree height, $H_{\text{max}}$ (m) | Initial observed | Baseline annual precipitation | 0.85*** |
|  |  | Baseline annual VPD | -0.67** |
|  |  | Baseline summer VPD | -0.78*** |
| Change in $H_{\text{mean}}$ ($\text{m yr}^{-1}$) | Observed | Change in annual precipitation | 0.50* |
|  | Allometry | Change in annual precipitation | 0.66** |
| Relative change in $H_{\text{mean}}$ ($\% \text{ yr}^{-1}$) | Observed | Change in annual precipitation | 0.57* |
|  | Allometry | Change in annual precipitation | 0.52* |
| Change in $H_{\text{max}}$ ($\text{m yr}^{-1}$) | Observed | Change in annual precipitation | 0.48* |
|  | Allometry | Change in annual precipitation | 0.57* |
| Relative change in $H_{\text{max}}$ ($\% \text{ yr}^{-1}$) | Observed | Change in annual precipitation | 0.61** |
|  | Demography | Change in annual precipitation | 0.53* |
|  | Allometry | Change in annual precipitation | 0.46* |

### Species-level allometric shifts

For individual species within ecoregions, most allometric shifts were towards greater height for a given DBH (Fig. 4 and Figs. S14-S15). For canopy trees in the East, 68% (263/385) of allometric shifts were significant (*p* < 0.05), with 95% (250/263) of significant shifts being positive (greater height for a given DBH; Fig. 4a and Fig. S14a). For canopy trees in the West, 50% (128/257) of allometric shifts were significant, with 68% (87/128) of significant shifts being positive (Fig. 4b and Fig. S14b). Allometric shifts for understory trees also tended to be positive (Fig. S15): in the East, 54% (167/311) were significant (96% positive); and in the West, 32% (34/107) were significant (74% positive). When compared at a common reference DBH for a given species, allometric shifts for understory trees were larger, on average, than for canopy trees (Table 3). Understory and canopy allometric shifts were positively correlated across species × ecoregion pairs, with stronger correlations in the West than the East (Table 3).

**Fig. 4.**
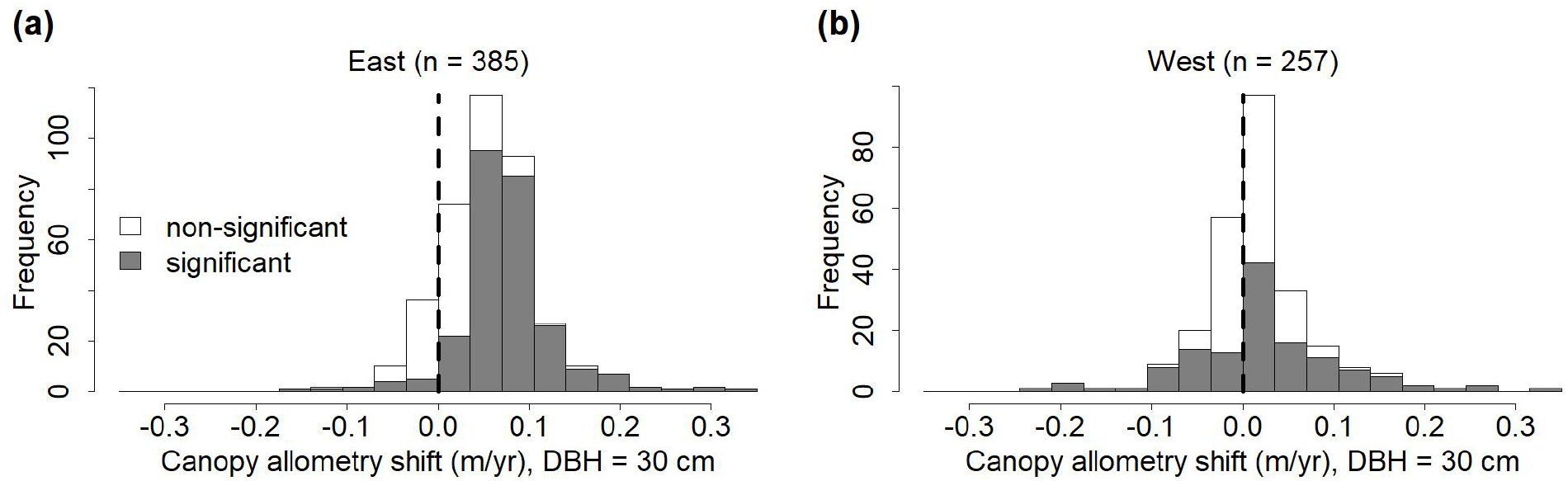
Allometric shifts for canopy trees with a diameter at breast height (DBH) of 30 cm in the **(a)** eastern and **(b)** western U.S. for the no-management dataset (non-plantation inventory plots with no reported management activities during the study period). The sample size above each histogram is the number of allometric models (species within ecoregions). Of the 263 significant (*p* < 0.05) shifts in the East, 250 (95%) were positive (shift towards taller trees for a given DBH). Of the 128 significant shifts in the West, 87 (68%) were positive. For a tree of DBH *x*, the allometric shift (m yr^−1^) is *αx*/(*k* + *x*), where *α* is the rate of change of the asymptotic height, and *k* is the DBH at which half of the asymptotic height is attained (Eqs. 2-3). Allometric shifts for canopy and understory trees with 10 cm DBH are shown in Figs. S14-S15.

**Table 3.**
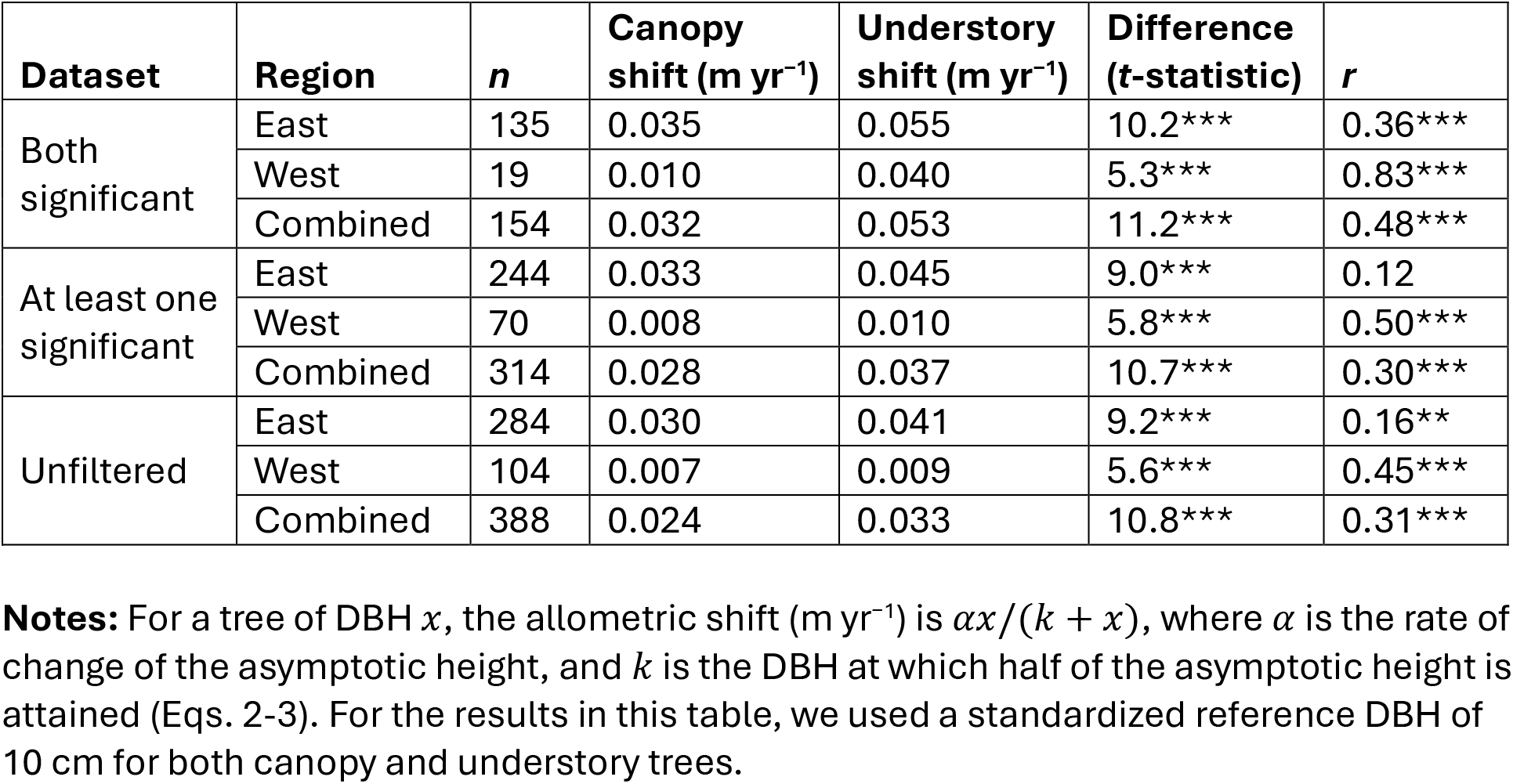
Comparisons of allometric shifts between canopy and understory trees in the eastern U.S., the western U.S., and both regions combined. “Dataset” describes filtering based on whether the temporal shift (*α* in Eq. 2) was significant for both members of a pair (canopy and understory trees of a given species × ecoregion), significant for at least one of the pair, or unfiltered (all canopy-understory pairs). The sample size (*n*) is the number of canopy-understory pairs. “Canopy shift” and “Understory shift” columns report allometric shifts for trees with a diameter at breast height (DBH) of 10 cm, averaged across species × ecoregions. The “Difference” column reports a variance-weighted paired *t*-test comparing canopy and understory means (understory shifts were greater, on average, in all cases). Pearson’s *r* is the correlation between canopy and understory allometric shifts across pairs (*r* > 0 indicates that larger canopy shifts tend to be associated with larger understory shifts). Significance levels: ** *p* < 0.01; *** *p* < 0.001.

## Discussion

Changes in forest canopy height from were largely positive in the eastern U.S. (i.e., canopy height increased) but varied in sign in the western U.S., mirroring patterns in aboveground wood biomass (AGB; Fig. 1 and Fig. S5) and recent changes in forest carbon stocks (Domke et al., 2026). Our analysis synthesizes data from around 60,000 inventory plots in non-plantation forests that were measured and remeasured at different times. Our study period roughly spans from 2003 (2.5^th^ percentile of initial plot measurement date) to 2025 (97.5^th^ percentile of final plot measurement date), with mean initial and final measurement dates of roughly 2009 and 2020, respectively. Climate change does not follow a smooth linear progression (Alley et al., 2003), and our study may not fully capture changes in U.S. forests over recent decades. For example, the current megadrought in the western U.S. began abruptly in 2000, with no clear trend in regional mean soil moisture from 2000-2021 (Williams et al., 2022). Our study would not capture changes in AGB or canopy height caused by the initial onset of this megadrought. Nevertheless, our study provides a systematic, broad-scale view of recent changes in AGB, canopy height, and its demographic and allometric components. The tight coupling between AGB and canopy height, and the weak coupling between AGB and tree density (Figs. S6-S7), highlights the importance of canopy height for understanding and monitoring forest biomass (Chave et al., 2019; Jucker et al., 2017; Lefsky et al., 2002), a large component of forest carbon storage (Pan et al., 2024).

Our study provides several lines of support for the hydraulic limitation (HL) hypothesis, which posits that tree heights are limited by hydraulic stress (Koch et al., 2004; McDowell & Allen, 2015; Moles et al., 2009; Ryan et al., 2006; Ryan & Yoder, 1997; Tao et al., 2016). According to the HL hypothesis, canopy height should increase across space and time with increasing moisture, and allometric shifts for understory trees should be more positive (or less negative) than for canopy trees, assuming the latter experience greater hydraulic stress (Myers et al., 1987; Niinemets, 2007). Canopy height increased across U.S. ecoregions with moisture (increasing precipitation and decreasing VPD; Table 2), consistent with the HL hypothesis and previous global studies (Tao et al., 2016; J. Zhang et al., 2016). Similarly, observed height change and its allometric component (Δ*H*_allom_; Table 1) increased with annual precipitation (more positive height change in ecoregions with more positive precipitation change; Table 2). Non-significant correlations between height change and other hydrological variables (trends in summer precipitation change, summer VPD, and annual VPD) may reflect the failure of our simplistic and coarse-scale climate analysis to capture the complex soil hydrology of the mountainous western U.S. (Hamlet et al., 2007; Harpold & Molotch, 2015). Regardless of the reason for the non-significant correlations, all significant correlations that we did observe were consistent with the HL hypothesis. Finally, consistent with the HL hypothesis, understory trees tended to have more positive allometric shifts than canopy trees (Table 3).

Allometric shifts tended to be positive (increasing canopy height for a given DBH), albeit highly variable, at both species (Figs. 4 and S14-S15) and ecoregion levels (Figs. 3 and S10-S11). Beyond the modest positive correlations between Δ*H*_allom_and precipitation change (Table 2), the widespread positive allometric shifts across climatically diverse regions (Figs. S12-S13) suggest a simple, common explanation, such as increasing atmospheric CO_2_. This explanation is supported by allometric shifts observed for *Pinus taeda* in a free-air CO_2_ enrichment experiment (Kim et al., 2020), as expected from CO_2_-driven increases in water-use efficiency (Keenan et al., 2013; van der Sleen et al., 2015). Thus, high CO_2_ may mitigate the hydraulic stress that trees would otherwise experience due to rising VPD in some ecoregions. This suggests a modification to simple predictions derived from the HL hypothesis. For example, the simple prediction that tree height should decrease with rising VPD could be revised as: changes in tree height should be negatively correlated with changes in VPD. This revision allows for the possibility of widespread increases in tree height (due to rising CO_2_), but with smaller height increases in regions with larger VPD increases.

Compared to Δ*H*_allom_, the demography component of canopy height change (Δ*H*_dem_; Table 1) was more variable (Figs. 3 and S10) and more strongly correlated with observed height change (Fig. 2). Δ*H*_dem_, which includes both shifts in DBH distributions and shifts in abundance between short- and tall-allometry species, was negative in much of the western U.S. (Figs. 3 and S10). Intensifying fire, insect, and drought disturbances in this region (Anderegg et al., 2013; Weed et al., 2013; Westerling et al., 2006) are expected to reduce forest canopy heights directly through reduced mean forest age, and also indirectly as taller-statured species are replaced by shorter ones (McDowell et al., 2020; McDowell & Allen, 2015). These direct and indirect effects may have both contributed to negative Δ*H*_dem_ in the western U.S. Despite negative Δ*H*_dem_in most western ecoregions, canopy height did not systematically decline across the western U.S. because negative Δ*H*_dem_was roughly balanced by allometric shifts towards greater height (positive Δ*H*_allom_).

Intraspecific shifts in allometry have important implications for monitoring forest canopy height and biomass. First, applying standard (static-in-time) height-DBH allometries to western U.S. forests over our study period (∼2003-2025) would lead to the erroneous conclusion that canopy heights decreased throughout most of this region. In contrast, our results suggest that allometric shifts roughly compensated for the decreases in western U.S. canopy heights that would have occurred due to demographic change alone. Second, if allometries (on average) are shifting over time towards greater height for a given DBH, as suggested by our results, ignoring these shifts will lead to biased estimates of biomass change. If biomass is monitored by combining remote-sensing-derived height change with static biomass-height relationships, then biomass change will be overestimated (because in reality, biomass for a given height will decrease as allometries shift taller). Conversely, if biomass is monitored by combining DBH measurements with static biomass-DBH allometries that ignore height, then biomass change will be underestimated (because in reality, biomass for a given DBH will increase as allometries shift taller). The FIA program, which conducts the nationwide forest inventory of the U.S., uses biomass allometries that depend on measurements of both DBH and height (Westfall et al., 2024), which in principle would capture the effects of changing allometries. Some other biomass allometries also incorporate both DBH and height (e.g., Chave et al., 2014), but many ground-based studies rely on DBH-only biomass allometries due to the labor costs of measuring tree heights, particularly in tall, dense forests.

How large is the bias associated with ignoring allometric shifts? Because biomass is roughly proportional to height (Chave et al., 2014; Domke et al., 2012), it is easy to estimate the magnitude of the potential bias. For example, consider a 30-cm-DBH tree in a population whose height allometry is shifting by 0.1 m yr^−1^ (a large but not extreme shift in our dataset; Fig. 4). Assuming a 30-cm-DBH tree is roughly 20 m tall (Figs. S2-S3), a 0.1 m yr^−1^ allometric shift implies a 0.5% yr^−1^ increase in height and biomass (e.g., an increase in the height of 30-cm-DBH trees from 20 m to 21 m over one decade, and a proportional increase in biomass). This effect size is similar to the theoretical limit of tree growth responses to CO_2_ fertilization (Walker et al., 2021); i.e. a proportional response to a ∼0.5% yr^−1^ increase in atmospheric CO_2_ since 1960. Thus, the effect size of allometric shifts on biomass may be of a similar magnitude as other important global change drivers.

Given the importance of accurately monitoring forest canopy height and biomass, we recommend the following: (*i*) assessments of canopy height change should be based on measured heights or dynamic allometries that account for shifts over time; (*ii*) shifts over time in biomass-height relationships should be quantified and accounted for in remote-sensing-based biomass monitoring; and (*iii*) ground-based biomass monitoring should account for allometric shifts either by measuring both DBH and height (and using biomass allometries that are sensitive to both) or by measuring enough heights to account for allometric shifts when estimating biomass.

## Acknowledgements

This work was funded by USDA Forest Service grant 21-JV-11242305-097.

