## Supplemental Information for "Forest canopy height change due to demographic and allometric shifts"

**Contents:**

Table S1

Figures S1-S15

**Table S1.** Sample sizes for plots, canopy trees, and understory trees at initial ($t$ = 0) and final ($t$ = $T$) plot measurements for (1) all non-plantation plots and (2) non-plantation plots with no reported management activities during the study period ("no-management dataset"). "Non-plantation" plots are classified by FIA as "natural stands" (as opposed to artificial regeneration) but may include harvest or other management during the study period. Ecoregion codes are defined in Fig. S1. Plots: number of remeasured FIA plots. $Can_0$ and $Can_T$: number of live canopy trees (open-grown, dominant, or codominant FIA crown class) with a measured DBH and height at $t$ = 0 ($Can_0$) and $t$ = $T$ ($Can_T$). $Und_0$ and $Und_T$: number of live understory trees (overtopped FIA crown class) with a measured DBH and height at $t$ = 0 ($Und_0$) and $t$ = $T$ ($Und_T$). Tree sample sizes indicate the number of tally trees used to estimate canopy and understory allometries, rather than tree densities (see table notes).

| | **All non-plantation plots** | | | | | **No-management dataset** | | | | |
|---|---|---|---|---|---|---|---|---|---|---|
| **Eco-region** | **Plots** | **$Can_0$** | **$Can_T$** | **$Und_0$** | **$Und_T$** | **Plots** | **$Can_0$** | **$Can_T$** | **$Und_0$** | **$Und_T$** |
| 210 | 10,427 | 196,752 | 216,383 | 35,496 | 92,624 | 8,931 | [illegible] | [illegible] | [illegible] | [illegible] |
| 220 | 7,318 | 103,238 | 97,840 | 39,026 | 63,484 | 6,173 | [illegible] | [illegible] | [illegible] | [illegible] |
| 230 | 9,885 | 161,542 | 162,439 | 83,454 | 93,996 | 8,347 | [illegible] | [illegible] | [illegible] | [illegible] |
| 240 | 107 | 2,310 | 2,245 | 539 | 729 | 91 | [illegible] | [illegible] | [illegible] | [illegible] |
| 250 | 1,730 | 22,932 | 24,920 | 4,968 | 7,539 | 1,491 | [illegible] | [illegible] | [illegible] | [illegible] |
| 260 | 268 | 8,316 | 8,344 | 1,631 | 1,833 | 219 | [illegible] | [illegible] | [illegible] | [illegible] |
| 310 | 4,244 | 56,712 | 57,041 | 1,778 | 2,124 | 3,737 | [illegible] | [illegible] | [illegible] | [illegible] |
| 320 | 1,180 | 11,412 | 10,942 | 167 | 200 | 1,074 | [illegible] | [illegible] | [illegible] | [illegible] |
| 330 | 771 | 10,454 | 10,060 | 755 | 1,044 | 695 | [illegible] | [illegible] | [illegible] | [illegible] |
| 340 | 2,051 | 34,298 | 32,809 | 679 | 670 | 1,965 | [illegible] | [illegible] | [illegible] | [illegible] |
| 410 | 62 | 1,177 | 1,284 | 261 | 177 | 56 | [illegible] | [illegible] | [illegible] | [illegible] |
| M210 | 2,726 | 56,275 | 63,954 | 12,719 | 36,890 | 2,398 | [illegible] | [illegible] | [illegible] | [illegible] |
| M220 | 3,380 | 52,281 | 50,664 | 28,494 | 37,045 | 3,089 | [illegible] | [illegible] | [illegible] | [illegible] |
| M230 | 538 | 9,986 | 10,829 | 3,903 | 5,196 | 482 | [illegible] | [illegible] | [illegible] | [illegible] |
| M240 | 4,805 | 107,950 | 106,905 | 20,549 | 21,031 | 4,287 | [illegible] | [illegible] | [illegible] | [illegible] |
| M260 | 3,309 | 72,678 | 66,236 | 8,073 | 8,495 | 2,924 | [illegible] | [illegible] | [illegible] | [illegible] |
| M310 | 1,045 | 18,766 | 16,479 | 663 | 758 | 977 | [illegible] | [illegible] | [illegible] | [illegible] |
| M330 | 11,477 | 253,987 | 229,035 | 16,616 | 17,230 | 10,451 | [illegible] | [illegible] | [illegible] | [illegible] |
| M340 | 1,447 | 29,993 | 29,365 | 795 | 754 | 1,404 | [illegible] | [illegible] | [illegible] | [illegible] |
| **Total** | **66,770** | **1,211,059** | **1,197,774** | **260,566** | **391,819** | **58,791** | [illegible] | [illegible] | [illegible] | [illegible] |

**Notes:** Numbers of canopy and understory trees do not reflect relative densities, because FIA uses different sample areas for trees in different DBH classes. Furthermore, sample sizes in the table are for trees with a measured DBH and height, which is smaller than the total number of tallied trees due to missing height data. Large increases in understory tree numbers from $t$ = 0 to $t$ = $T$ (e.g., ecoregions 210, 220, and M210) are due to missing height data at $t$ = 0 rather than changes in tree density (Figs. S6-S7).

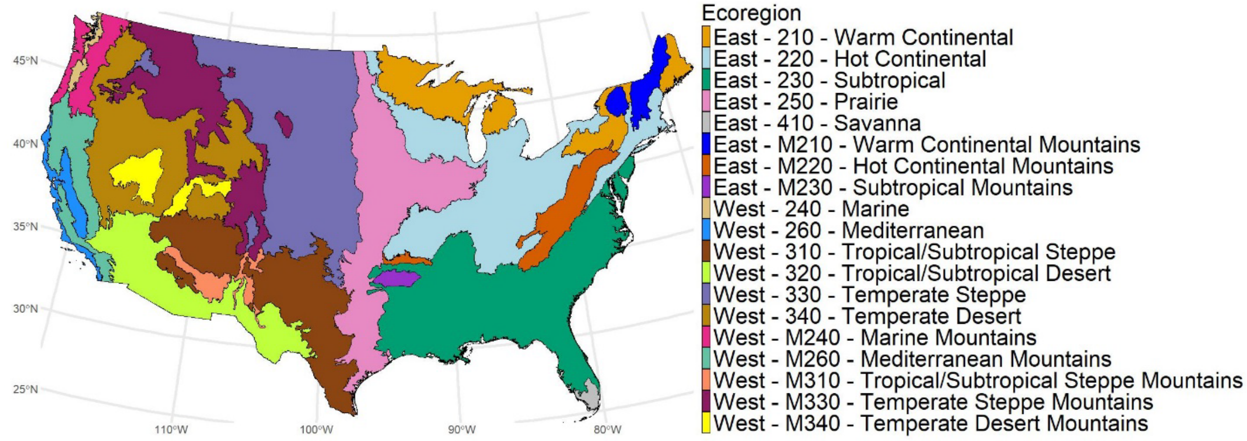


**Fig. S1.** Map of ecoregions used in this study. These ecoregions are the division-level units of McNab et al. (2007) and represent regions of relatively similar climate, bedrock, and potential vegetation. Note that some ecoregions are geographically discontinuous.

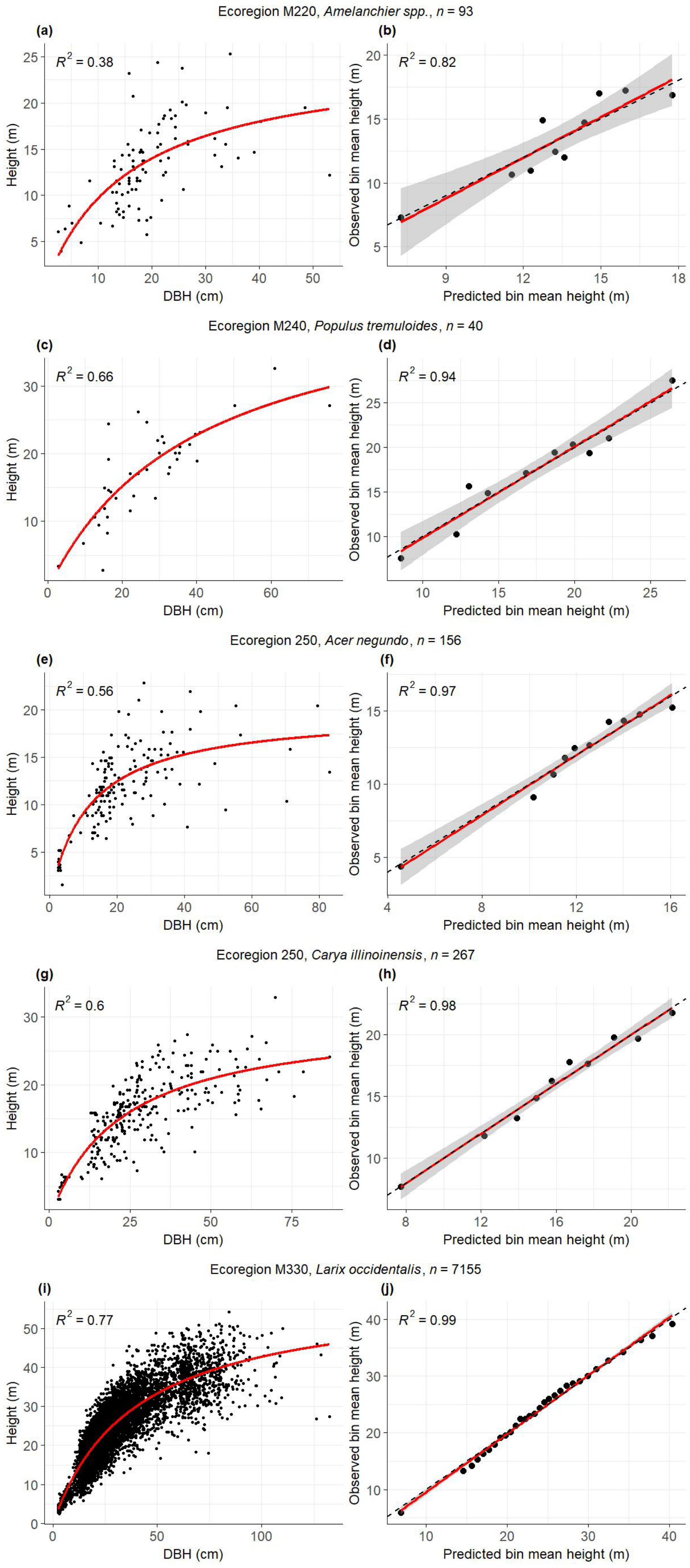


**Fig. S2.** Example diagnostic plots for static allometry models (Eq. 1) for canopy trees. **Left column:** Fitted allometries and raw data ($t$ = 0 data). **Right column:** Observed vs. predicted bin-level mean height (red solid line and gray shading: regression of observed vs. predicted with 95% confidence band; black dashed line is 1:1 line). The examples shown are the 10th, 30th, 50th, 70th, and 90th percentiles of the bin-level $R^2$ values ($R^2_{\mathrm{bin}}$ in Methods) across the 642 species × ecoregion static models for canopy trees.

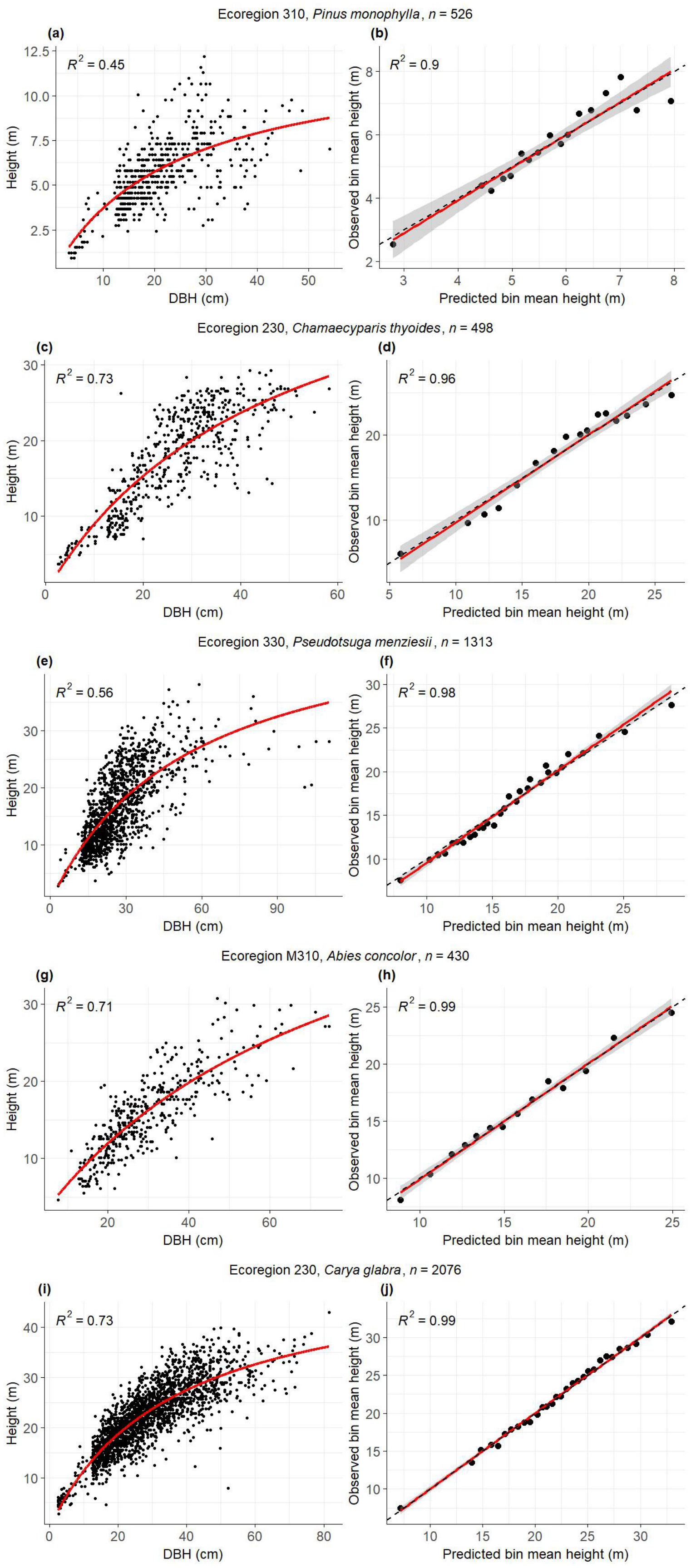


**Fig. S3.** Example diagnostic plots for dynamic allometry models (Eq. 2) for canopy trees. **Left column:** Fitted allometries and raw data ($t$ = 0 and $t$ = $T$ data combined). The $R^2$ values are based on the measurement date ($t$) of each observation, whereas fitted curves are plotted at the mean $t$ across all $t$ = 0 and $t$ = $T$ data for a given species × ecoregion. Thus, allometric shifts over time contribute to the visual noise in the left column but are accounted for in the $R^2$ calculations. **Right column:** Observed vs. predicted bin-level mean height (red solid line and gray shading: regression of observed vs. predicted with 95% confidence band; black dashed line is 1:1 line). The examples shown are the 10th, 30th, 50th, 70th, and 90th percentiles of the bin-level $R^2$ values ($R^2_{\mathrm{bin}}$ in Methods) across the 642 species × ecoregion dynamic models for canopy trees.

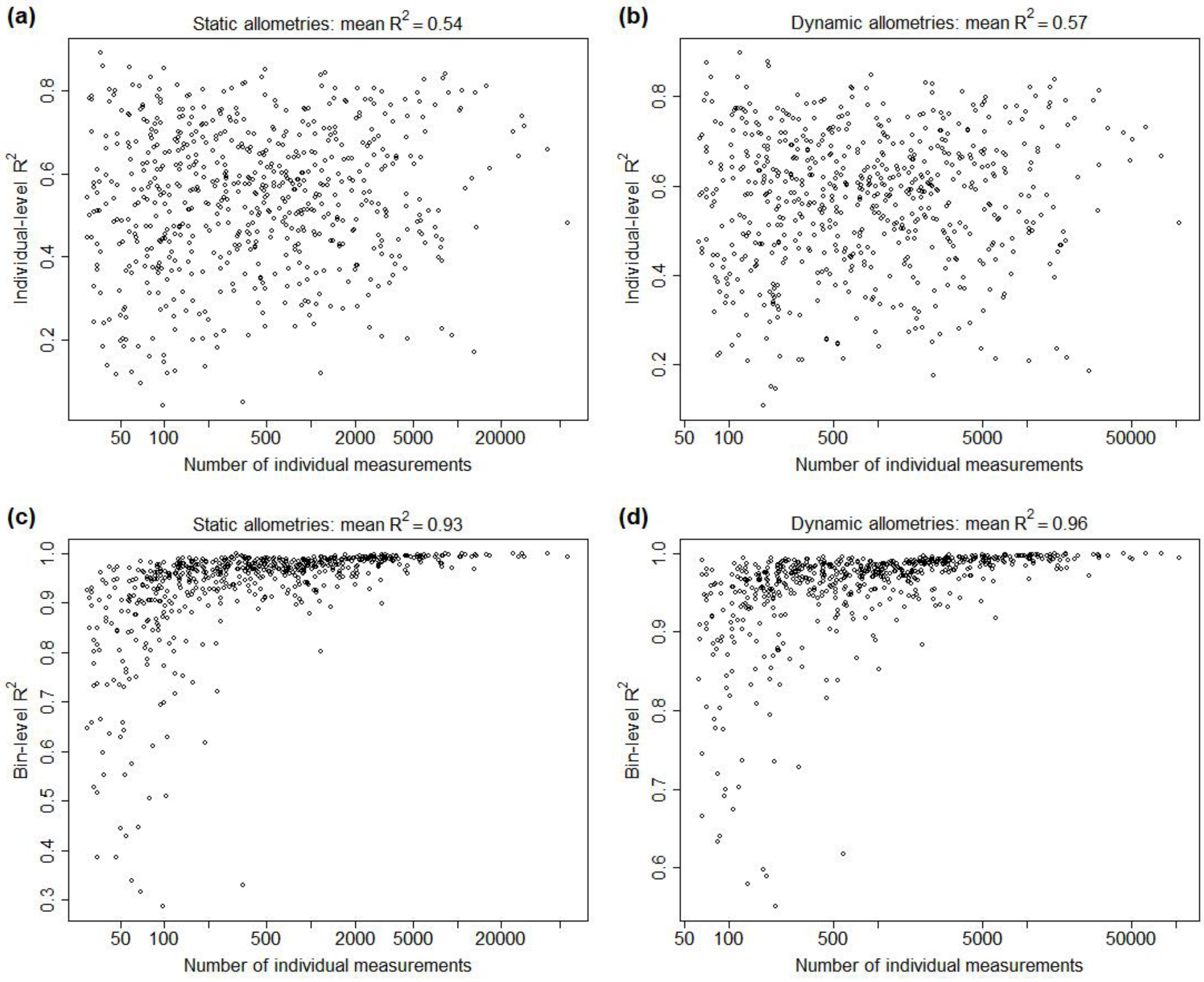


**Fig. S4.** Relationships between $R^2$ and sample size at the individual tree level **(a-b)** and the aggregated bin level **(c-d)** for canopy-tree allometries. Individual-level $R^2$ values indicate how much tree-level variation is explained by the height-DBH relationship, and bin-level $R^2$ values indicate how well the functional form fits the data, after averaging over individual-level noise. Titles above each panel indicate the type of allometry (static or dynamic) and the mean $R^2$ across the 642 canopy allometries (species within ecoregions) represented in each panel. $R^2$ values for understory allometries (418 species × ecoregion combinations) were similar (not shown).

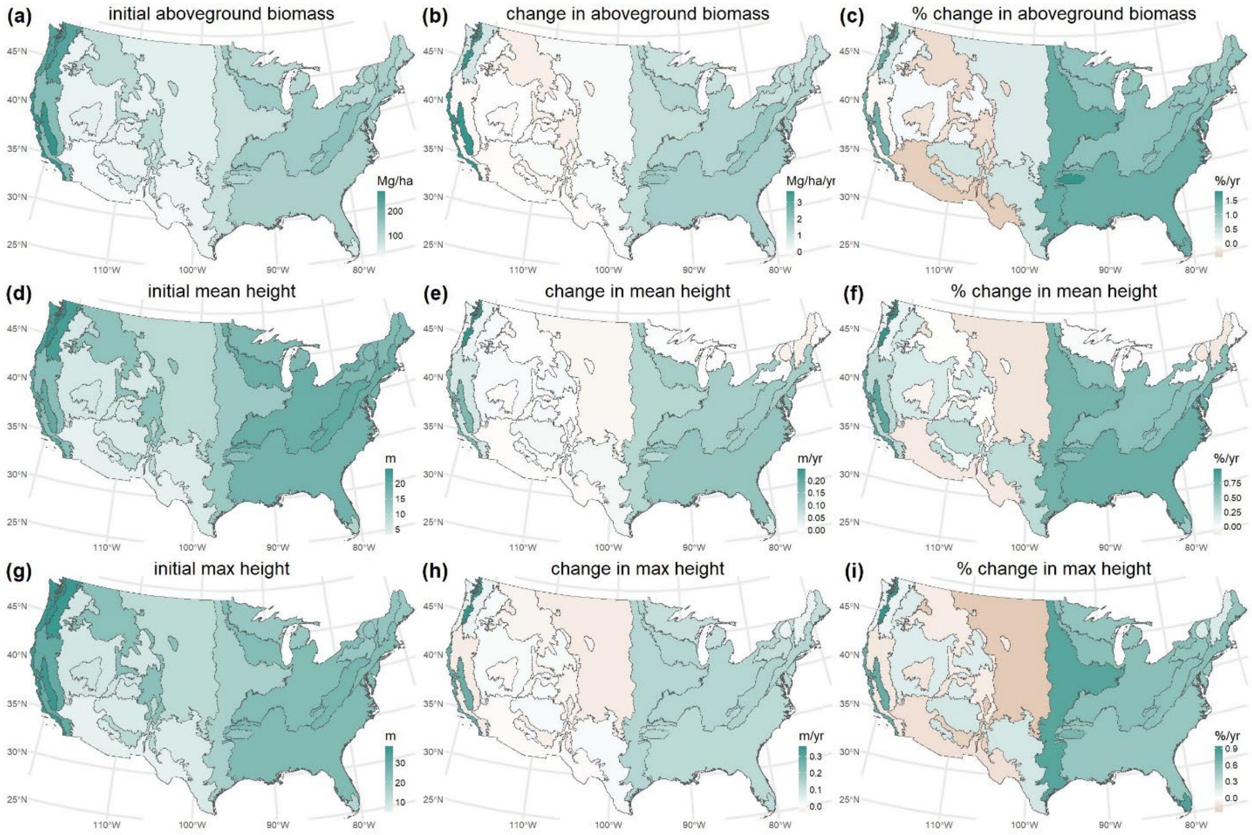


**Fig. S5.** Changes in aboveground biomass **(a-c)**, mean canopy height **(d-f)**, and maximum tree height **(g-i)** for all non-planation inventory plots, including those with harvest or other management activities during the study period. Colors indicate the mean value across inventory plots in each ecoregion (Fig. S1). **Left column:** ecoregion means for the initial measurement of each plot (mean initial date ~2009). **Middle column:** mean change from the initial to final measurement of each plot (mean final date ~2020). **Right column:** relative change = 100 × (middle column)/(left column).

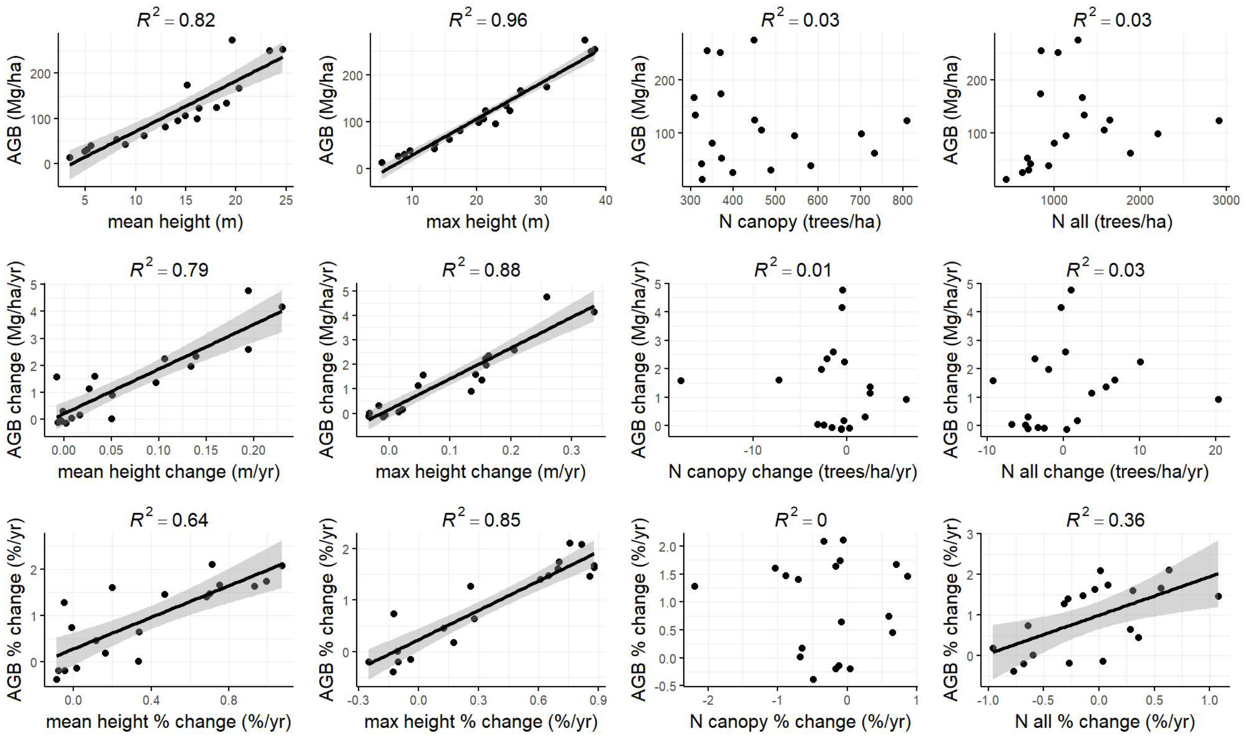


**Fig. S6.** Relationships between aboveground biomass (AGB) and four metrics of forest structure—mean canopy height, maximum tree height, canopy tree density (“N canopy”), and total tree density (“N all”)—across U.S. ecoregions (*n* = 19) for the no-management dataset (non-plantation inventory plots with no reported management activities during the study period). **Top row:** ecoregion means for the initial measurement of each forest inventory plot (mean initial date ~2009). **Middle row:** mean change from the initial to final measurement of each plot (mean final date ~2020). **Bottom row:** relative change = 100 × (middle row)/(top row). Regression lines are shown with shaded 95% confidence intervals for significant relationships ($p < 0.05$).

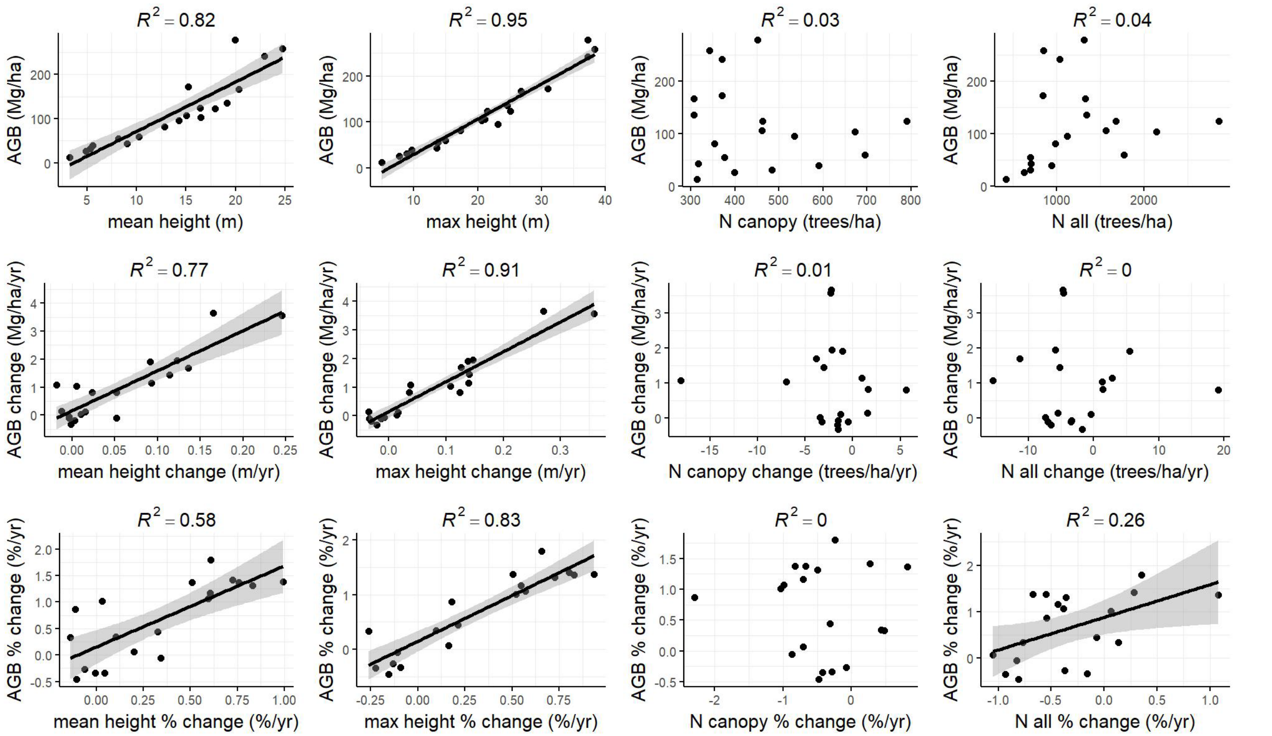


**Fig. S7.** Relationships between aboveground biomass (AGB) and four metrics of forest structure for all non-plantation inventory plots, including those with harvest or other management activities. Other details are as in Fig. S6.

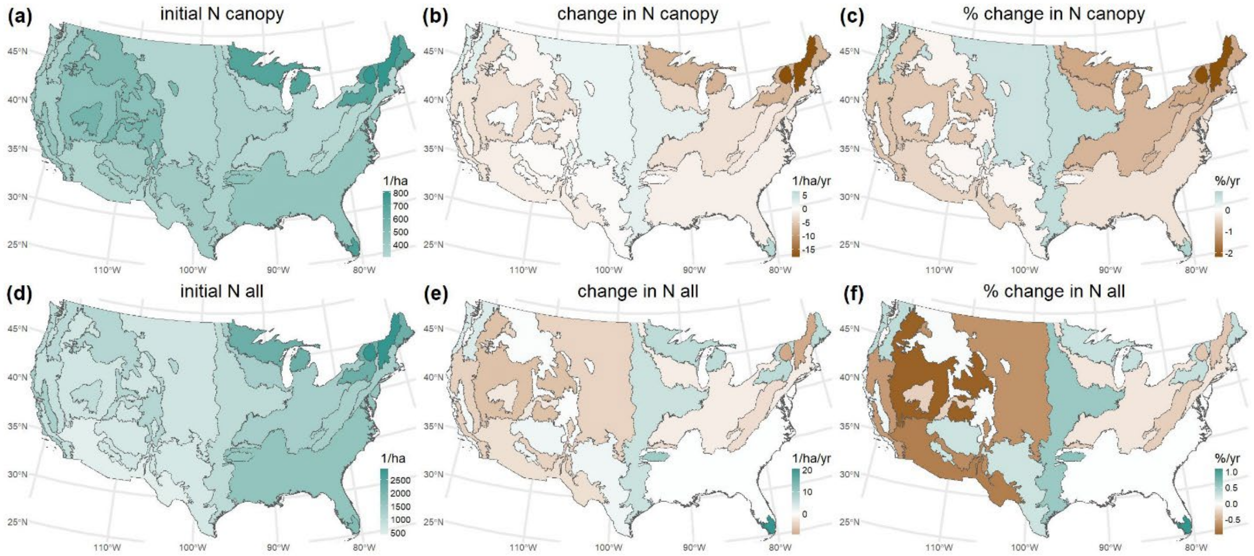


**Fig. S8.** Changes in the density (trees/ha) of canopy trees **(a-c)** and all trees **(d-f)** for the no-management dataset (non-plantation inventory plots with no reported management activities during the study period). “Canopy trees” includes trees with an open-grown, dominant, or codominant FIA crown class. “All trees” includes trees with DBH ≥ 2.54 cm. Colors indicate the mean value across inventory plots in each ecoregion (Fig. S1). **Left column:** ecoregion means for the initial measurement of each plot (mean initial date ~2009). **Middle column:** mean change from the initial to final measurement of each plot (mean final date ~2020). **Right column:** relative change = 100 × (middle column)/(left column).

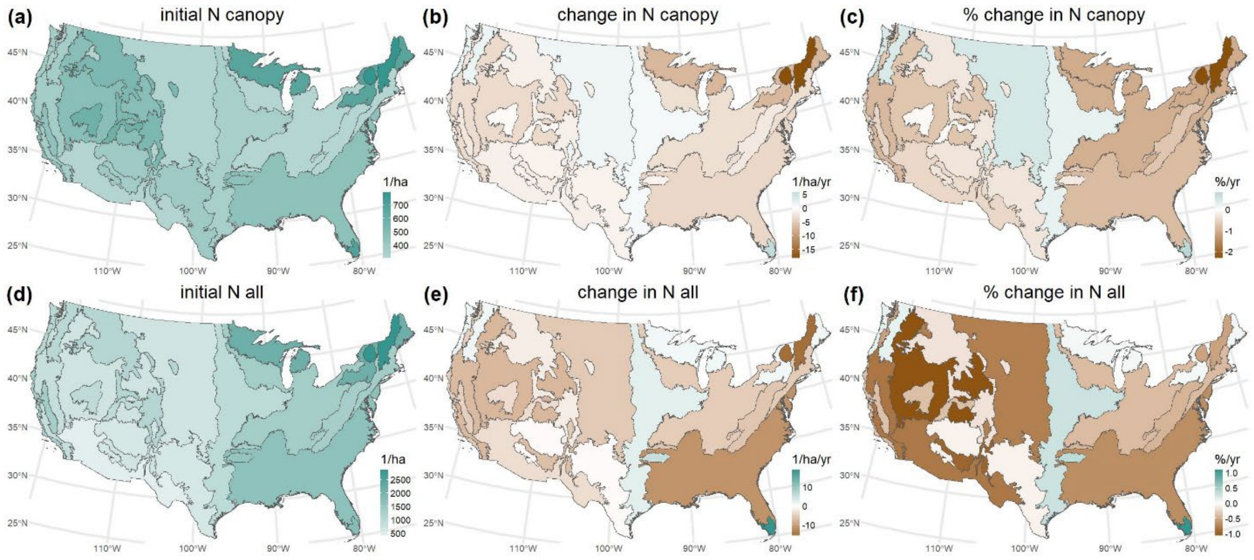


**Fig. S9.** Changes in the density (trees/ha) of canopy trees **(a-c)** and all trees **(d-f)** for all non-planation inventory plots, including those with harvest or other management activities during the study period. Other details are as in Fig. S8.

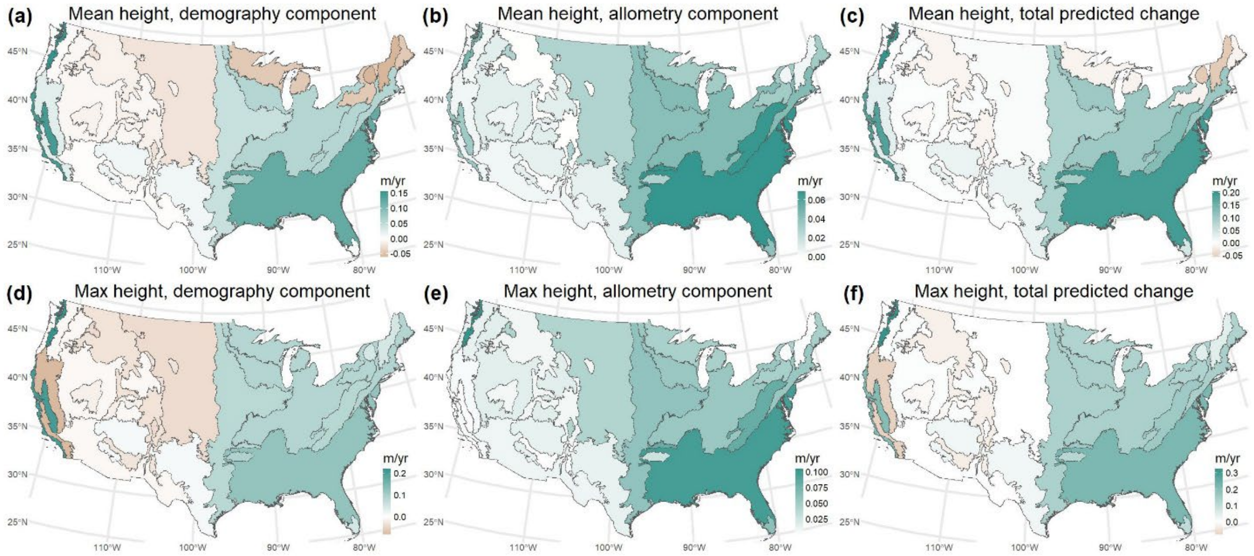


**Fig. S10.** Predicted change (m/yr) in mean canopy height **(a-c)** and maximum tree height **(d-f)** for the no-management dataset (non-plantation inventory plots with no reported management activities during the study period). Details are as in Fig. 3, except for units (m/yr here; %/yr in Fig. 3).

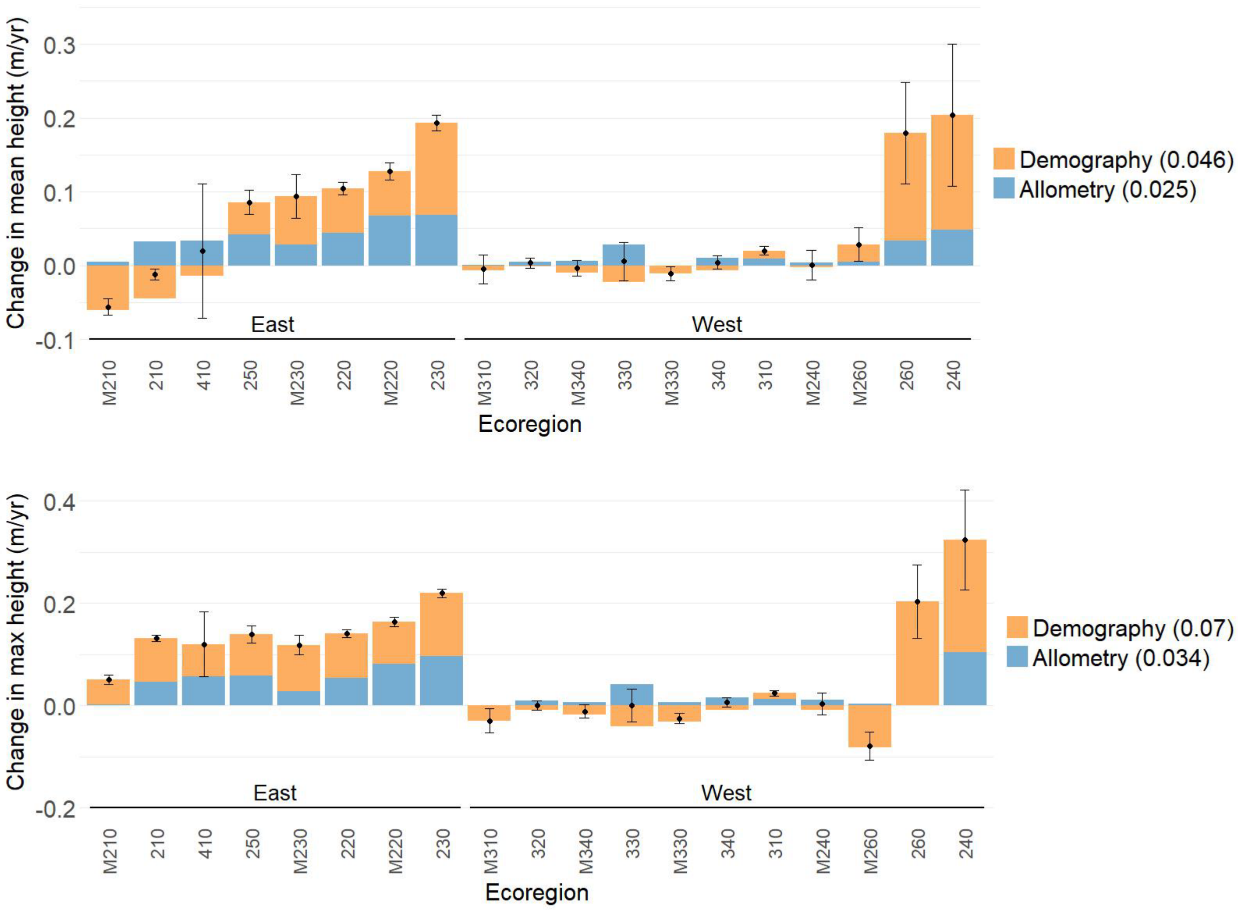


**Fig. S11.** Demography and allometry components of height change by ecoregion (**top:** mean canopy height; **bottom:** maximum tree height) for the no-management dataset (non-plantation inventory plots with no reported management activities during the study period). The demography component represents height change due to shifts in both the DBH distribution and species composition, holding intraspecific allometries constant. The allometry component represents height change due to intraspecific shifts in allometry, holding the DBH distribution and species composition constant. Numbers in the legend are means across ecoregions of the absolute value of each component. Black symbols are the total predicted change (sum of components) with 95% confidence intervals. Within East and West regions, ecoregions are ordered by increasing observed change in mean canopy height.

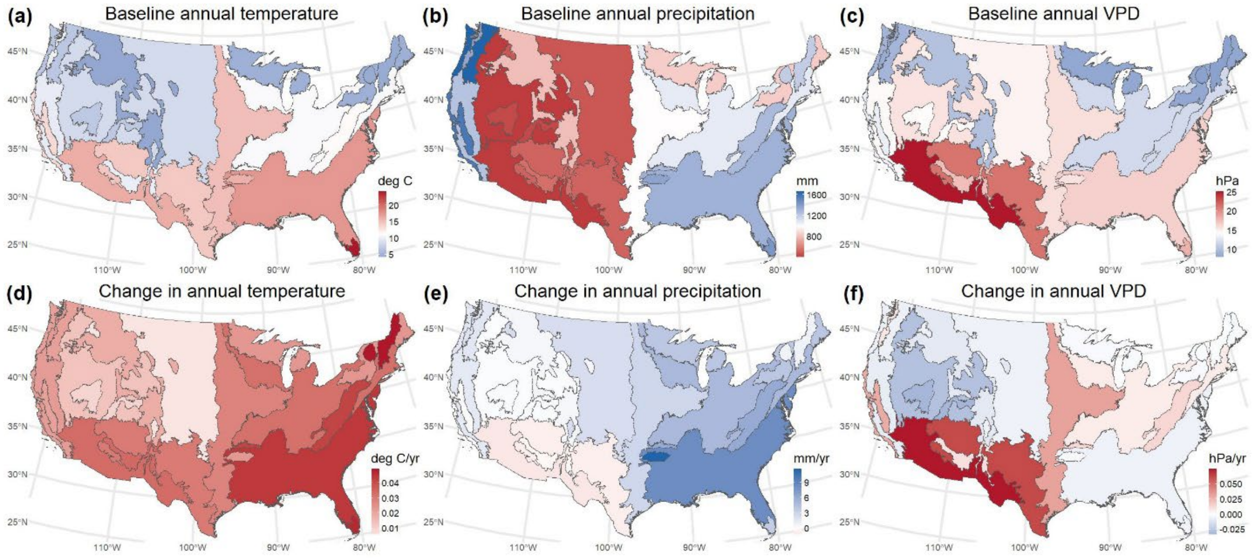


**Fig. S12.** Baseline means from 1990-2002 **(a-c)** and trends from 2003-2025 **(d-f)** for annual temperature, precipitation, and VPD. Colors indicate the mean value across inventory plots in each ecoregion (Fig. S1). Color scales are oriented so that red corresponds to hot/dry, and blue corresponds to cool/wet.

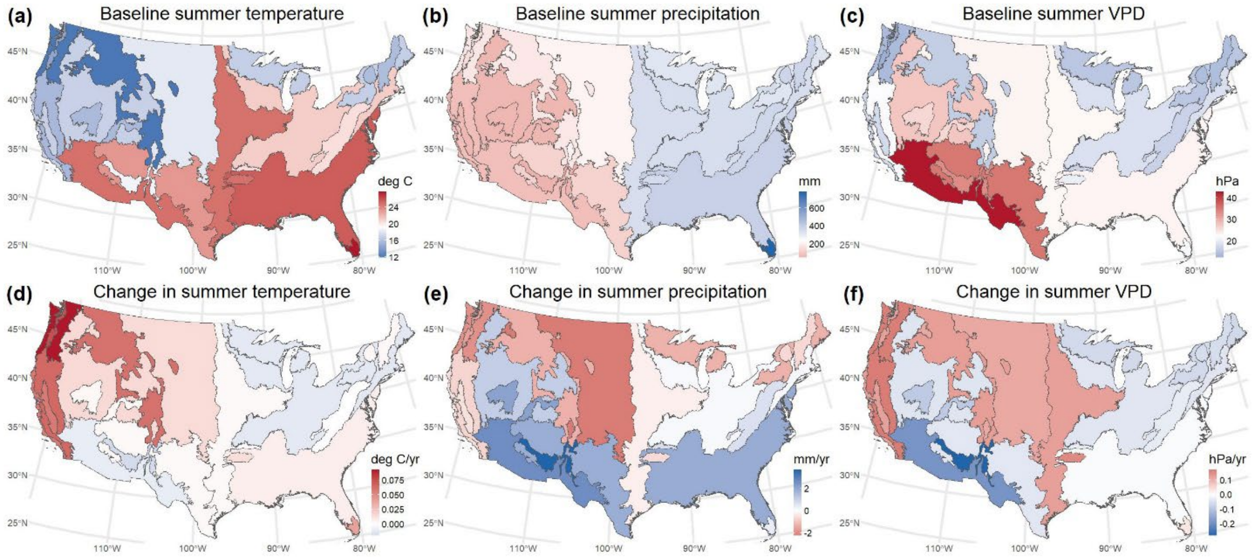


**Fig. S13.** Baseline means from 1990-2002 **(a-c)** and trends from 2003-2025 **(d-f)** for summer (June, July, August) temperature, precipitation, and VPD. Colors indicate the mean value across inventory plots in each ecoregion (Fig. S1). Color scales are oriented so that red corresponds to hot/dry, and blue corresponds to cool/wet.

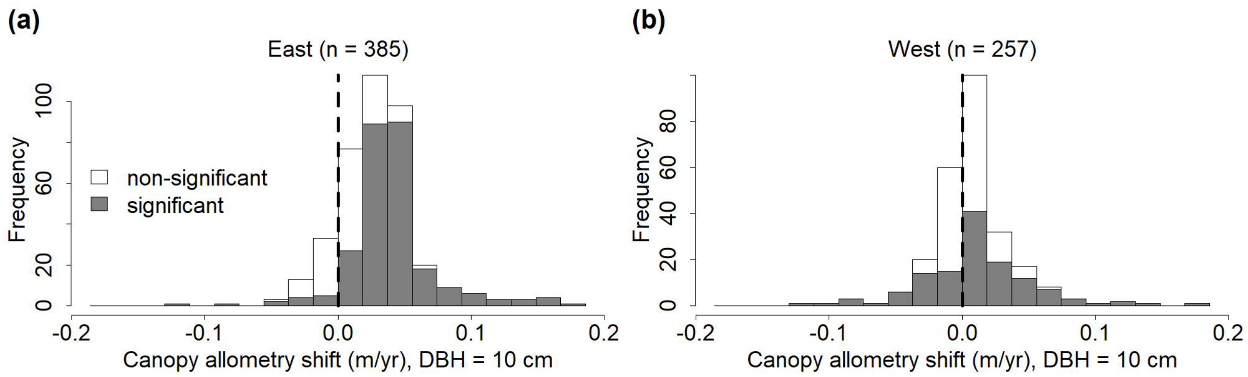


**Fig. S14.** Allometric shifts for canopy trees with a diameter at breast height (DBH) of 10 cm in the **(a)** eastern and **(b)** western U.S. for the no-management dataset (non-plantation inventory plots with no reported management activities during the study period). Details are as in Fig. 4. Statistical significance of shifts is the same as for 30 cm DBH trees (Fig. 4), but the magnitude of each shift depends on DBH. Here, canopy trees with 10 cm DBH are evaluated for comparison with 10 cm DBH understory trees (Fig. S15; Table 3).

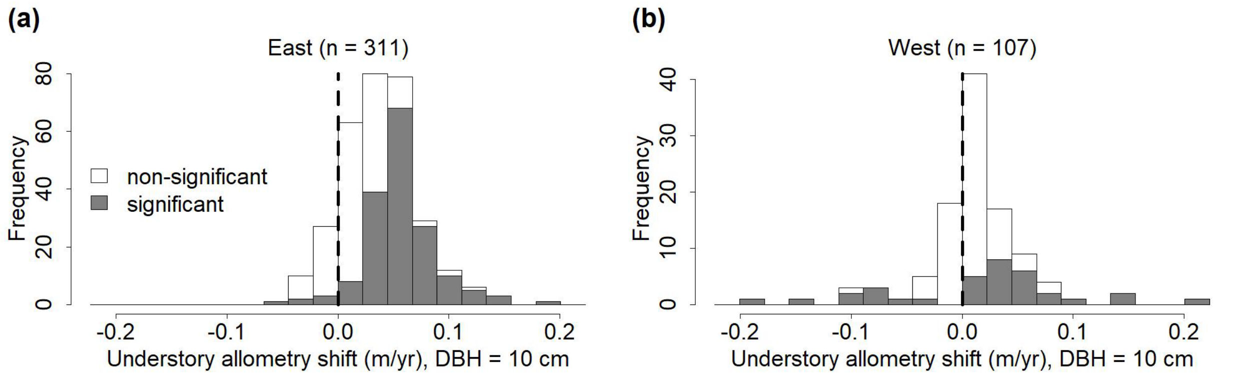


**Fig. S15.** Allometric shifts for understory trees with a diameter at breast height (DBH) of 10 cm in the **(a)** eastern and **(b)** western U.S. for the no-management dataset (non-plantation inventory plots with no reported management activities during the study period). The sample size above each histogram is the number of allometric models (species within ecoregions). Of the 167 significant ($p < 0.05$) shifts in the East, 161 (96%) are positive (shift towards taller trees for a given DBH). Of the 34 significant shifts in the West, 25 (74%) are positive. Other details are as in Fig. 4.
